# E-cadherin-dependent phosphorylation of EGFR governs a homeostatic feedback loop controlling intercellular junction viscosity and collective migration modes

**DOI:** 10.1101/2023.12.04.570034

**Authors:** Chaoyu Fu, Florian Dilasser, Shao-Zhen Lin, Marc Karnat, Aditya Arora, Harini Rajendiran, Hui Ting Ong, Nai Mui Hoon Brenda, Sound Wai Phow, Tsuyoshi Hirashima, Michael Sheetz, Jean-François Rupprecht, Sham Tlili, Virgile Viasnoff

**Affiliations:** Mechanobiology Institute, National University of Singapore, 5a engineering drive 1 117411 Singapore; Aix Marseille Univ, Université de Toulon, CNRS, CPT (UMR 7332), Turing Centre for Living systems, Marseille, France; Department of Biomedical Engineering, National University of Singapore, 4 Engineering Drive 3, 117583, Singapore; Aix Marseille Univ, IBDM (UMR 7288), Turing Centre for Living systems, Marseille, France; CNRS IRL 3639, 5a Engineering drive 1, 117411 Singapore

## Abstract

Actomyosin tension has been shown to be a ubiquitous driver of tissue morphogenesis^1, 2^. The Rho pathway, a prominent regulatory network influencing cortical tension, plays a central role in both tissue reorganisation and cell migration^3–6^. While viscous dissipation in the actin network is commonly regarded as a constant passive parameter in cell migration in both 2D and 3D contexts, there is limited knowledge concerning the regulation of dissipative forces arising from viscous drag between cells during collective rearrangement. Here, we found that the phosphorylation of Epithelial Growth Factor Receptor (EGFR) downstream of *de novo* E-cadherin adhesion^7, 8^ orchestrates a feedback loop, thereby governing intercellular viscosity via the Rac pathway regulating actin dynamics. Our findings highlight how the E-cadherin-dependent EGFR activity controls the migration mode of collective cell movements independently of intercellular tension. Combining molecular cell biology, micropatterning, and *in silico* simulation, our work suggests the existence of a regulatory loop by which cells can tune junctional actin viscosity, with implications for the phenomenology of morphogenetic movements.

## Main

We postulated the existence of a feedback loop between cell junction elongation and E-cadherin-dependent phosphorylation of EGFR at junctions. This hypothesis was tested on Madin-Darby Canine Kidney (MDCK) cells with 2D migration in a culture dish. All experiments were conducted using both serum-free and serum-rich media, with consistently similar phenotypes observed, albeit more pronounced effects in serum-free conditions. All presented results pertain to serum-free conditions (**Methods**).

We first compared the migration of MDCK cells under control conditions and following selective inhibition of EGFR phosphorylation using Erlotinib at 1μM. Quantification of cell movement on 2D or 1D line patterns (**Extended Data Fig. 1a, c**) demonstrated that the migration of single isolated cells remained insensitive to EGFR inhibition. Consequently, we ruled out the possibility that EGFR inhibition directly altered cell-substrate interactions as well as their single cell migration potential. In contrast, EGFR inhibition significantly impacted collective cell migration on 2D sparse islets and 1D line patterns (**Fig. 1a, Extended Data Fig. 1b, d, and Supplementary Video 1)**. In the former case, EGFR inhibition markedly reduced the cellular swirling motion of cells by inhibiting EGFR phosphorylation. The velocity at which each contact changes length during migration (**Methods**), exhibited a 2-fold decrease upon inhibition (**Fig. 1b**). Thus, EGFR dephosphorylation reduced the dynamics of cell junction deformation.

**Fig 1:**
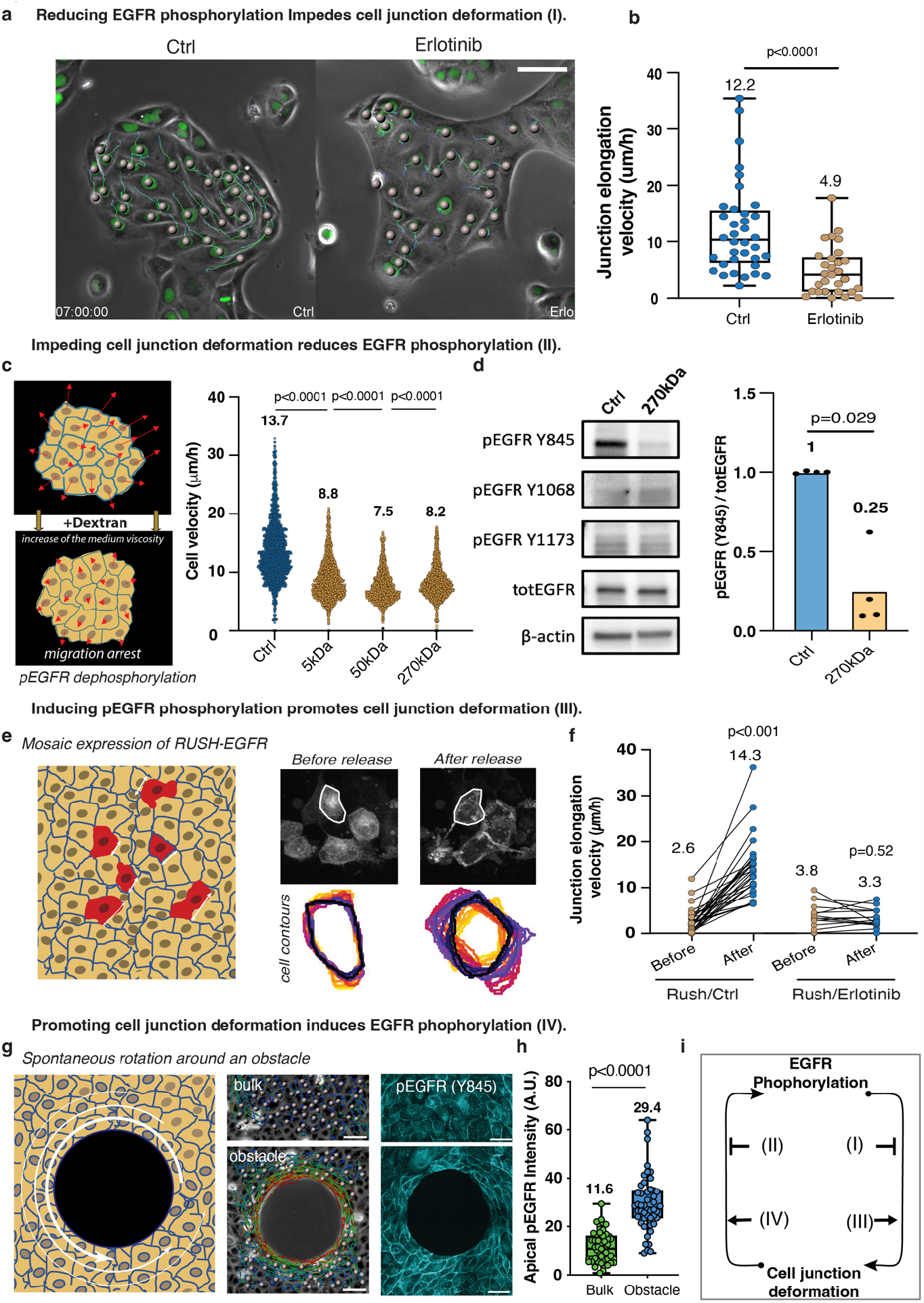
A positive feedback loop between apical EGFR phosphorylation and cell junction deformation. **a**.Representative patches of MDCK cells under control and EGFR-inhibited conditions (Erlotinib at 1μM) including the tracking or individual cell trajectories. Scale bar: 100 μm. **b**.Quantification of individual junction elongation velocities in the patches (mean value ± s.d.) n_Ctrl_ = 36 junctions and n_Erlotinib_ = 29 junctions from 3 independent experiments, two-tailed unpaired t-test, p < 0.0001. **c**.Schematics of the experiment for cell arrest by dextran addition. Quantification of individual cell migration velocity 10 minutes after adding dextran with various molecular weights (mean value ± s.d. n=1735-1925 cells from 3 independent experiments.) **d**.Western Blot and its quantification of EGFR phosphorylated states (Y845) before and after cell arrest from 4 independent experiments, two-tailed unpaired t-test, p = 0.029. **e**.Experimental setup schematics (left) and segmented contours quantification (right) of cell mosaically expressing RUSH-EGFR before and after its release from the endoplasmic reticulum. **f**.Quantifications of junction elongation velocities upon the release of EGFR, under control and pEGFR-inhibited conditions. n_Rush/Ctrl_ = 28 junctions and n_Rush/Erlotinib_ = 15 junctions from 3 independent experiments, two-tailed paired t-test, p_Rush/Ctrl_ < 0.001, p_Rush/Erlotinib_ = 0.52. **g**.Schematics of the physical induction of cell elongation around obstacles (left). Images of cells encircling obstacles and in bulk regions including single cell tracking and apical localization of pEGFR-Y845 by immunostaining. Scale bar: 50 μm. **h**.Quantifications of apical pEGFR-Y845 intensity around obstacles (mean value ± s.d. n_Bulk_ = 50 junctions and n_Obstacle_ = 48 junctions from 3 independent experiments, two-tailed unpaired t-test, p<0.0001.) **i**.Diagram of a positive feedback loop between apical EGFR phosphorylation and cell junction deformation.

Reciprocally, we impeded the physical deformation of contacts during migration by supplementing the medium with 50 μg/mL of dextran (5, 50 and 270 kDa) to increase the medium viscosity. We carefully ensured that the addition of dextran induced minimal osmotic shock (**Extended Data Fig. 2a**). In media with higher viscosity (270kDa dextran), the migration speed of individual isolated cells decreased by 28 % (**Extended Data Fig. 2b)**. Within cohesive patches consisting of 20 to 50 cells, both collective migration and contact deformation ceased within 10 minutes of adding 50 μg/mL dextran (**Fig. 1c, Extended Data Fig. 2c and Supplementary Video 2**). The individual cell velocity within the patch decreased from 13.7 ± 0.13 μm/h (N=1735 cells) in the control group to 8.2 ± 0.06 μm/h (N=1925) for 270kDa dextran. Immunostaining revealed a significant reduction in the phosphorylation of EGFR at the apical junction (**Extended Data Fig. 2d**). Western blots confirmed a 75% reduction in the phosphorylation of the Src-dependent site Y845 pEGFR^9^, while the other phosphorylation sites (Y1068, Y1173) remained inactive under serum-free conditions (**Fig. 1d**). Our data strongly suggest that the physical arrest of cell junction deformation directly or indirectly leads to the dephosphorylation of apical pEGFR at its Src-dependent site.

We subsequently investigated whether enhancing pEGFR would favour the dynamics of contact deformation. To do so, we transfected MDCK cells with EGFR coupled to the RUSH system (**Methods**). The RUSH-EGFR construct was sequestered on the ER membrane until released by biotin addition in the culture medium^10^. We established a high-confluence, non-polarised (95%) MDCK monolayer, resulting in a mosaic expression of RUSH-EGFR, with small patches of positive cells amid non-expressing control MDCK cells (**Fig. 1e**). Prior to the addition of biotin, the cells displayed limited junctional localization and weak recruitment of apical EGFR (**Supplementary Video 3**). Upon the addition of biotin, the RUSH-positive cells showed a substantial recruitment of EGFR at the cell contacts, leading to a 5-fold increase (from 2.6 ± 0.5 to 14.3 ± 1.2 μm/h) in their junction elongation velocity, quantified using Cellpose neural network segmentation of the cell contours (**Fig. 1e, f and Methods**). In RUSH-positive cells, EGFR localized exclusively to cell-cell contacts within minutes (**Extended Data Fig. 3a**). To validate these findings, we repeated the experiment in the presence of Erlotinib (1μM). Despite the relocalization of EGFR to the junctions (**Extended Data Fig. 3b and Supplementary Video3**), the junction elongation velocity remained as low as the control, with very limited cellular rearrangements. These results strongly suggest that the burst increase in pEGFR at cell-cell contacts favours the dynamics of contact deformation.

Conversely, we induced the physical elongation of cell junctions by embedding obstacles (non-adhesive disks with a diameter of 200μm, **Methods**) into high confluence monolayers (**Fig. 1g**). Only the limited number of cell layers that spontaneously elongated and encircled the obstacles displayed deforming junctions (**Fig. 1g and Supplementary Video 4**) and elevated levels of apical pEGFR (**Fig. 1g**), in sharp contrast to the immobile bulk cells (**Fig. 1h**). A treatment with Erlotinib inhibited the elongation and circumrotation of the cells around the obstacle (**Supplementary Video 4**). Taken together, our findings support the hypothesis of a positive feedback loop (**Fig. 1i**) between apical EGFR phosphorylation and junction elongation. We subsequently delved into the molecular mechanisms underlying this phenomenon.

The absence of the soluble ligand EGF and the specific phosphorylation of Y845 suggested an E-cadherin (Ecad)-dependent activation of EGFR^7, 8, 11^. Confluent patches of WT-MDCK cells displayed a two-fold decrease in apical pEGFR compared to sub-confluent patches (27.6 ± 11.1 A.U. vs 51.6 ± 19.0 A.U.) (**Fig. 2a**). In contrast, Ecad-KO tissues showed consistently low levels of apical pEGFR in both confluent and sub-confluent cases (4.0 ± 2.3 A.U. vs 6.1 ± 3.2 A.U.), while still forming cohesive patches (likely due to K-cadherins, quantified in (**Extended Data Fig. 4c**) with a proper junctional actin structure. In Ecad-KO MDCK cells with rescued expression of Ecad (Ecad-Res), the pEGFR levels in confluent and sub-confluent tissues returned to their control values (22.2 ± 4.0 A.U. vs 30.2 ± 7.9 A.U.). Furthermore, when EGFR was inhibited by adding Erlotinib (1 μM), the apical pEGFR for both confluent and sub-confluent patches in WT-MDCK, Ecad-KO MDCK, and Ecad-Res MDCK all dropped to low levels (**Extended Data Fig. 4a, b**).

**Fig 2:**
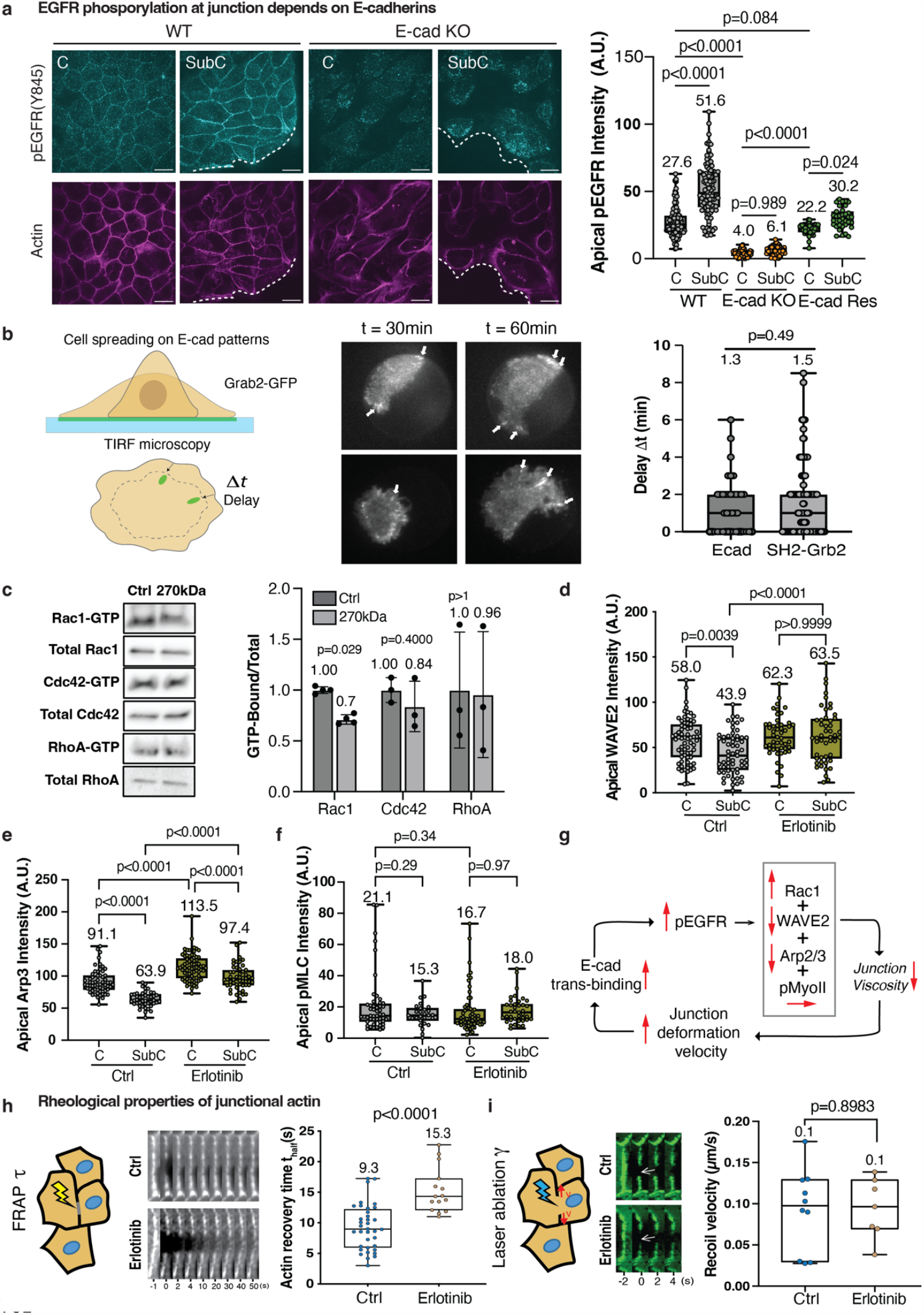
E-cadherin-dependent phosphorylation of EGFR fine-tunes actin dynamics with minimal impact on cortical tension. **a**.Immunostaining of apical pEGFR (Y845) and actin in wild-type (WT) and E-cadherin knock-out (Ecad-KO) MDCKs on confluent (C, left) and sub-confluent (SubC, right) regions. Scale Bar: 20 μm. Quantification of apical pEGFR in WT, Ecad-KO and Ecad-KO-rescued (Ecad-Res) tissues. (WT: n_C_ = 122 junctions, n_SubC_ = 106 junctions from 4 independent experiments, p < 0.0001; Ecad-KO: n_C_ = 67 junctions, n_SubC_ = 61 junctions from 3 independent experiments, p = 0.9893; Ecad-Res: n_C_ = 47 junctions, n_SubC_ = 34 cell junctions from 3 independent experiments, p = 0.0239. Ordinary one-way ANOVA Tukey’s test. **b**.Schematic (left) and time-lapse imaging (middle) of SH2-Grb2 (tdEOS) and E-cadherin (GFP) localization during cell spreading on E-cadherin-coated patterns. Quantification of the recruitment speed of E-cadherin and SH2-Grb2 (right). n_Ecad_ = 40 cell adhesions and n_SH2-Grb2_ = 132 cell adhesions from 3 independent experiments, two-tailed unpaired t-test, p=0.49. **c**.Pull-down assays on Rho family GTPases (Rac1, Cdc42 and RhoA) and quantification of GTP-bound GTPases post cell arrest by dextran. n_Rac1_ = 4 WB, p=0.029, n_Cdc42_ = 3 WB, p=0.4 and n_RhoA_ = 3 WB, p>1, two-tailed unpaired t-test. **d-f**. Quantification of apical WAVE2 (**d**), Arp3 (**e**) and pMLC (**f**) under control and EGFR inhibited conditions. Data are the mean value ± s.d. WAVE2: n_Ctrl, C_ = 68 cell junctions, n_Ctrl, SubC_ = 59 cell junctions from 2 independent experiments, n_Erlotinib, C_ = 50 cell junctions, n_Erlotinib, SubC_ = 48 cell junctions from 2 independent experiments; Arp3: n_Ctrl, C_ = 60 cell junctions, n_Ctrl, SubC_ = 43 cell junctions from 2 independent experiments, n_Erlotinib, C_ = 74 cell junctions, n_Erlotinib, SubC_ = 52 cell junctions from 2 independent experiments; pMLC: n_Ctrl, C_ = 49 cell junctions, n_Ctrl, SubC_ = 27 cell junctions from 2 independent experiments, n_Erlotinib, C_ = 62 cell junctions, n_Erlotinib, SubC_ = 39 cell junctions from 2 independent experiments. Ordinary one-way ANOVA Tukey’s test. **g**. Proposed model of E-cadherin-dependent phosphorylation of EGFR reducing junction viscosity through the regulation of Rac1, WAVE2, Arp2/3, fine-tuning actin dynamics, with minimal impact on cortical tension. **h-i**. Experimental schematics and characteristic images of fluorescence recovery after photobleaching (FRAP) (left) and laser ablation (right) experiments on intercellular junctions between GFP-Actin-MDCK cells, Scale Bar: 3 μm. Quantification of fluorescence recovery time (left) and recoil velocities (right) under control and pEGFR-inhibited conditions. FRAP: n_Ctrl_ = 36 cell junctions, n_Erlotinib_ = 15 cell junctions from 3 independent experiments, two-tailed unpaired t-test, p<0.0001; Laser ablation: n_Ctrl_ = 10 cell junctions, n_Erlotinib_ = 7 cell junctions from 2 independent experiments, two-tailed unpaired t-test, p=0.8983.

We further validated the direct phosphorylation of EGFR at adherens junctions. We used Total Internal Reflection Fluorescence (TIRF) microscopy to image the live recruitment of cytosolic SH2-Grb2 to the membrane as a proxy for EGFR phosphorylation^12^. MDCK cells stably expressing tdEOS-labelled SH2-Grb2 were left to spread on E-Cad coated circular patterns (25μm diameter) (**Fig. 2b and Methods**). SH2-Grb2 dynamically accumulated in elongated structures that dynamically followed the progression of cell edges with a time delay Δt. A parallel experiment using E-cad-GFP MDCK cells revealed a similar accumulation of E-cad in structures with similar time delay Δt (1.3 ± 0.2 min for E-cad-GFP, 1.5 ± 0.2 min for SH2-Grb2) (**Fig. 2b**). Our findings imply that the engagement of E-cad during junction elongation results in transient phosphorylation of EGFR. Consequently, the arrest of cell junction elongation leads to the dephosphorylation of pEGFR, whereas its physical induction promotes EGFR phosphorylation.

We then scrutinized the alteration of recruitment of actin regulators in the same conditions as above. EGFR phosphorylation is a major regulator of Erk, a kinase extensively implicated in collective cell migration mechanisms^13^. Monitoring Erk activity using Fluorescence Resonance Energy Transfer (FRET) did not reveal any changes following the addition of 50μg/mL dextran to migrating cells (**Extended Data Fig. 5a**). This suggests that Erk signaling is not downstream of EGFR in our experimental context. To further dissect the molecular events, we performed pulldown assays to gauge the activity of Rho family GTPases, which are key regulators of actin dynamics^14^. The introduction of dextran to migrating cells resulted in a 28.9% reduction in Rac1 activity, with no discernible effects on Cdc42 and RhoA (**Fig. 2c**). However, the broad nature of pulldown assays made it challenging to distinguish whether Rac1 activity was junctional or lamellipodial. Notably, Wave2, a downstream target of Rac1, was present at the apical side of the junction^15^ (**Extended Data Fig. 5b**). Surprisingly, the recruitment of Wave2 and Arp2/3, two factors promoting branched actin nucleation, was higher in confluent monolayers (58.0 ± 24.0 A.U. and 91.1 ± 18.8 A.U., respectively) than in sub-confluent patches (43.9 ± 22.9 A.U. and 63.9 ± 11.9 A.U., respectively), a difference that is substantially reduced upon treatment with Erlotinib (1μM) (**Fig. 2d, e and Extended Data Fig. 5b, c**). The amount of phosphorylated Myosin light chain (pMLC) remained unaffected by pEGFR both in confluent and sub-confluent culture conditions (**Fig. 2f and Extended Data Fig. 5d**). Our data advocate for a model wherein the trans binding of new E-cadherin in elongating junctions, increasing the junctional level of pEGFR, subsequently activating Rac1, decreasing levels of Wave2 and Arp2/3 with a constant myosin level, thereby establishing a balance between branched and linear junctional actin. **Fig. 2g** illustrates this feedback homeostatic loop.

Next, we evaluated the impact of EGFR phosphorylation on the turnover rate of junction actin using Fluorescence Recovery After Photobleaching (FRAP). We compared junctions in confluent monolayers, cells surrounding obstacles, and sub-confluent patches with or without EGFR inhibition (**Fig. 2h and Extended Data Fig. 6a-c**). In all cases, we observed a deceleration in actin turnover when pEGFR levels were lower. Additionally, we probed junctional tension by assessing fast actin recoil following laser ablation (**Fig. 2i**). We did not detect any substantial difference, proving that pEGFR does not regulate tension in this specific context. Finally, we probed the viscoelastic properties of junctions using Atomic Force Microscopy (AFM), revealing an increase in the loss modulus upon EGFR inhibition, with no changes in the elastic modulus (**Extended Data Fig. 6d-f**).

We previously reported that E-cad-dependent phosphorylation of EGFR in suspended cell doublets increases the velocity of *de-novo* junction formation^8^ and the toughness of their adhesion^16^. In all cases, the microscopic dynamics of the actin cortex are associated with a change in cell deformability, with minimal impact on cortical tension. This implies that the homeostasis of junction viscosity is regulated by the Ecad-dependent EGFR phosphorylation loop, effectively “self-lubricating” junction elongation.

By analogy with the transition from laminar to turbulent flows of fluids at various Reynolds numbers, we monitored how the inhibition of EGFR alters patterns of collective MDCK migration along fibronectin strips (width, 400 μm; length, 3000 μm) (**Methods**). **Fig. 3a** and **Supplementary Video 5** show the leading region (0-500 μm from the front) of migrating cells in ctrl and Erlotinib conditions over 12 hours. Although migration fronts collectively progressed at similar velocities in both conditions, inhibition of pEGFR abolished the vortices observed in the ctrl case, leading to more laminar flows with enhanced cellular elongation in the direction of migration. We subsequently quantified these qualitative observations. The high level of apical pEGFR in Ctrl was significantly reduced in the inhibitory case (**Fig. 3b**). In the bulk regions (3 mm away from the front), the level of apical pEGFR remained constant in both conditions. We computed the cellular flow lines in the monolayer (**Methods**) to established maps of cell velocities and flow vorticity (**Fig. 3c, d**). Additionally, we used CellPose to segment individual cells (**Methods**) and to quantify the individual level of strain on each cell. pEGFR inhibition resulted in around a 3-fold increase in cellular strain (0.06 ± 0.04 in Ctrl, 0.14 ± 0.06 in Erlotinib; n=1060 cells) (**Fig. 3e**). While the collective velocity of the migration fronts was not affected by pEGFR inhibition (15.0 ± 4.7 μm/h in Ctrl, 14.8 ± 2.9 μm/h in Erlotinib; N=16 strips), it substantially reduced the individual cell velocity within the monolayer (from 18.7 ± 2.0 μm/h to 12.6 ± 2.0 μm/h; n= 182 cells) (**Fig. 3g**). Likewise, the vorticity of the collective flow (**Fig. 3h)** was reduced by 2-fold (0.70 ± 0.10 h^-1^ to 0.33 ± 0.05 h^-1^; n= 182 cells), and the correlation length of the cell velocity increased by 2-folds (**Supplementary Fig. 4**).

**Fig 3:**
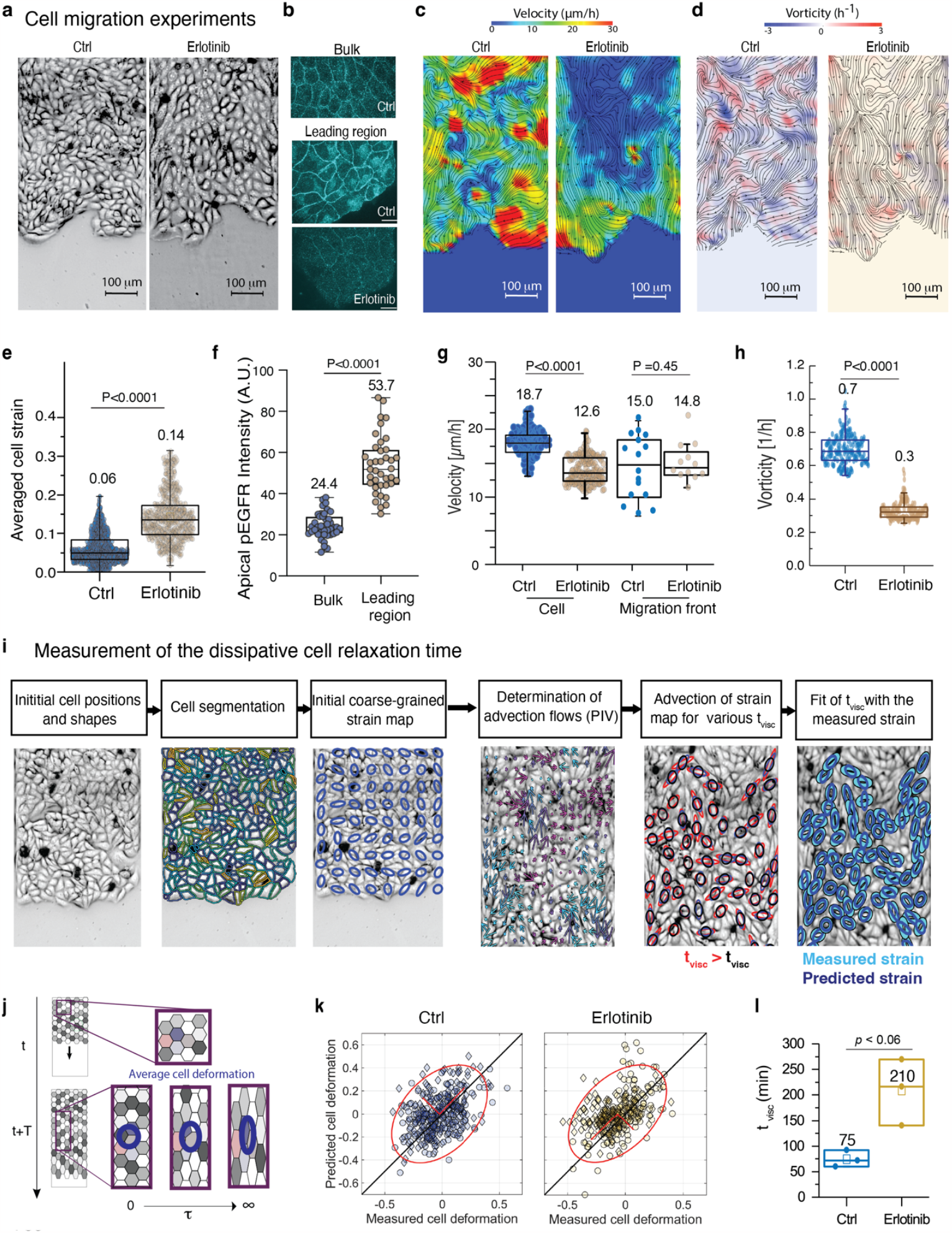
Phosphorylated EGFR (pEGFR) modulates cell deformability and influences collective migration. **a**.Phase-contrast images of MDCK monolayers migrating on fibronectin-coated line patterns under control and pEGFR-inhibited conditions (Erlotinib at 1 μM). Scale Bar: 100 μm. Five independent experiments yielded consistent results. **b**.Immunostaining of apical pEGFR(Y845) highlights its localization at cell junctions in bulk and leading regions under control and leading regions under pEGFR-inhibited conditions. Scale Bar: 20 μm. Four independent experiments corroborate these findings. **c-d**. Representative velocity (**c**) and vorticity (**d**) profiles with flow line maps, illustrate MDCK monolayer migration under control and pEGFR-inhibited conditions. Scale bar: 100 μm. Three independent experiments yielded consistent results. **e**.Quantification of cellular strain states in the monolayer under control and pEGFR-inhibited conditions. n_Ctrl_ = 1060 cells and n_Erlotinib_ = 1060 cells from 3 independent experiments, two-tailed unpaired t-test, p<0.0001. **f**.Quantification of apical localization of pEGFR (Y845) at the bulk and leading front region of the monolayer. n_Bulk_ = 42 cell junctions and n_leading_ = 39 cell junctions from 3 independent experiments, two-tailed unpaired t-test, p < 0.0001. **g**.Quantification of cell velocity (left) and migration front velocity (right) under control and pEGFR-inhibited conditions. n_ctrl, cell_ = 181 cells, n_Erlotinib, cell_ = 181 cells from 3 independent experiments, p<0.0001; n_ctrl, migration front_ = 16 strips, n_Erlotinib, migration front_ = 12 strips from 3 independent experiments, p=0.45, two-tailed unpaired t-test. **h**.Quantification of spatial correlation in the velocity field under control and pEGFR-inhibited conditions. n_ctrl_=181 cells, n_Erlotinib_=181 cells from 3 independent experiments, two-tailed unpaired t-test, p<0.0001. **i**.Schematic representation of the analysis pipeline to measure the average cell shape relaxation time in a migrating monolayer. Average cell strain profiles along the migration axis are depicted, with bold lines indicating mean values and narrow lines representing standard deviations. **j-k**. Correlative plots between measured and advection-based predicted cellular strain for the best fit of the viscoelastic time (t_visc_) under control and pEGFR-inhibited conditions. Two independent experiments yielded consistent results. **l**. Measured viscoelastic time (t_visc_) under control and pEGFR-inhibited conditions. n_ctrl_=3 strips, n_Erlotinib_=3 strips from 2 independent experiments, two-tailed unpaired t-test, p<0.06.

**Fig 4:**
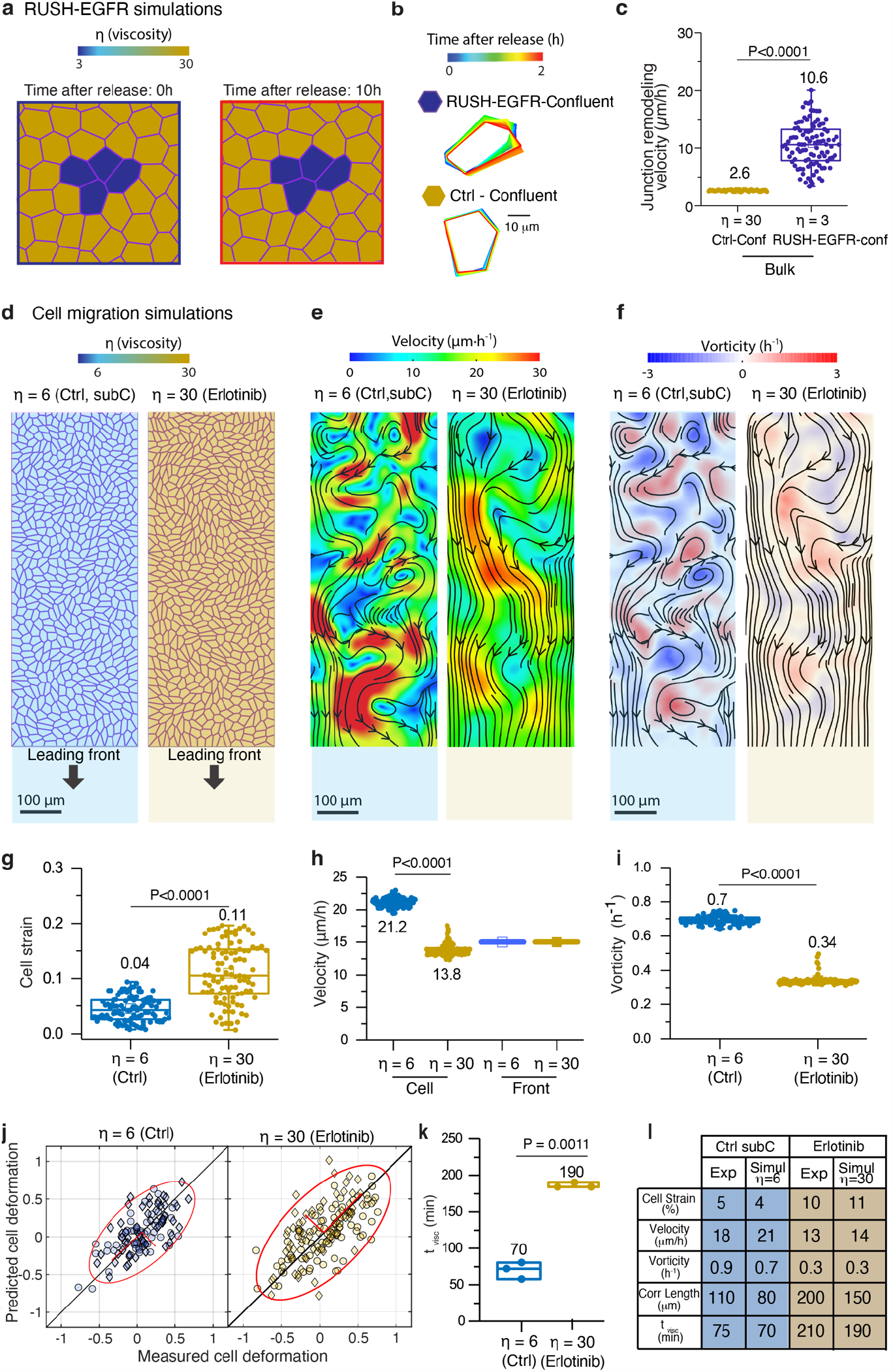
Modeling the impact of viscosity changes on epithelial migration. **a**.Simulated cellular arrangement in a vertex model with tension fluctuation, illustrating four low viscosity cells (dark blue, η=3) within a larger population of normal viscosity cells (orange, η=30). This models RUSH-EGFR activated cells within a large tissue of inactivated cells. **b**.Temporal variation of two representative cell profiles for low viscosity (RUSH-EGFR) cells (top) and control cells (bottom). **c**.Quantifications of the cell-cell junction remodeling velocity for control cells and low viscosity (RUSH-EGFR) cells (η=30 or η=3, respectively). Conf = Confluent. n_Ctrl_ = 101 cell junctions and n_RUSH-EGFR_ = 101 cell junctions from simulations, two-tailed unpaired t-test, p<0.0001. **d**.Simulated cellular arrangement in a vertex model with η=6 (light blue) and η=30 (orange), modelling the control and pEGFR-inhibited (Erlotinib) tissues, in the presence of cellular activity and an imposed uniform front migration speed. **e-f**. Velocity (**e**) and, vorticity fields (**f**), both with flow lines (black) for the model of control and pEGFR-inhibited conditions. **g-i**. Distribution in the simulated averaged cellular strain (**g**), velocity (displayed together with the imposed front migration speed) (**h**), and vorticity (**i**), under control and pEGFR-inhibited conditions (see **Methods** for averaging procedure). n_Ctrl, strain_ = 101 cells and n_Erlotinib, strain_ = 101 cells from simulations, two-tailed unpaired t-test, p<0.0001. n_Ctrl, velocity_ = 100 cells and n_Erlotinib, velocity_ = 100 cells from simulations, two-tailed unpaired t-test, p<0.0001. n_Ctrl, vorticity_ = 100 cells and n_Erlotinib, vorticity_ = 100 cells from simulations, two-tailed unpaired t-test, p<0.0001. **j**. Correlative plots between the measured and advection-based predicted cellular strain for the best fit of the viscoelastic time (t_visc_) under control (η=6) and pEGFR-inhibited (η=30) conditions. **k**. Viscoelastic time (t_visc_) for control (η=6) and pEGFR-inhibited (η=30). n_Ctrl_ = 3 strips and n_Erlotinib_ = 3 strips from simulations, two-tailed unpaired t-test, p=0.0011. **l**.Table summarizing the experimental and simulation results.

To further substantiate the hypothesis regarding a change in intercellular viscosity, we inferred the average shape relaxation time *t*_visc_ of cells within the monolayer. This parameter proved to be a reliable indicator of cellular viscoelasticity in migrating monolayers^17^. **Fig. 3i, j** illustrates the analysis procedure. Initially, we segmented phase-contrast images of the monolayer at a specific time point. Subsequently, we computed the average initial cell strain on a coarse-grained grid (**Methods**) and used an optic flow method to estimate the flow lines (**Methods**). The evolution of local cellular strain was then evaluated using the approach depicted in **Fig. 3i** and elaborated on in the **Supplementary Information**. The only free parameter in this equation is the intrinsic strain relaxation time t_visc_, which does not depend on the shear level experienced by the cells. By utilizing the strain map at t = 0 and solving the equation along the flow lines, we inferred the final strain map at a later time point (10 h). We varied t_visc_ to maximise the correlation between the observed and measured strain maps. The best fits lead to t_visc_ = 75 ± 15 min (R^2^=0.4±0.04; N=3) for the control group and t_visc_ = 210 ± 65 min (R^2^=0.34±0.2; N=3) for the pEGFR inhibition group (**Fig. 3k, l**). In the control conditions, cells exhibited shape relaxation when advected in swirling vortices, while under pEGFR inhibitory conditions, they elongated more in directed laminar, plug-like flows (**Supplementary Video 6)**. As pEGFR inhibition did not affect the single-cell migration (**Extended Data Fig. 1a, c**), the approximately 3-fold increase in cell shape relaxation times supports the hypothesis that the level of pEGFR controls viscous dissipation in cell-cell junction elongation.

Finally, we designed a vertex model with viscosity (**Supplementary Information**) to demonstrate that a modulation of intercellular viscosity can account quantitatively for our observations. In our model, the force balance at each tri-cellular junction is expressed as follows:

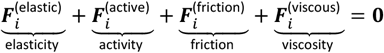

The first 3 terms are standard for vertex models^18, 19^ and correspond to: 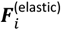 : the elastic forces accounting for the mechanical regulation of the cell shape^20–23^, that are assumed to derive from a mechanical energy *E* :

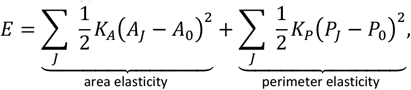

where the two terms account for the cell area elasticity and the cell perimeter elasticity, respectively. In detail, *K*_*A*_ and *K*_*P*_ are the area stiffness and perimeter stiffness of cells, respectively; *A*_*J*_ and *P*_*J*_ are the area and the perimeter of the *J*-th cell, respectively; *A*_0_ and *P*_0_ are the preferred area and preferred perimeter, respectively.

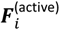 corresponding to the active fluctuations of the cortical tension, assumed to be a Gaussian white noise of amplitude Lambda (**Supplementary Information**), and

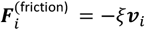

corresponding to the friction of the cells on the underlying substrates.

We complemented this description by adding a viscous dissipation term accounting for cortical deformation and cytoplasmic flows, which reads:

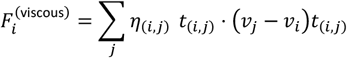

where *t*_(*i,j*)_ is a unit vector oriented between the vertex *i* and *j*, which is either another vertex, in which case *η*_(*i,j*)_ = *η*^(*s*)^ is the viscous modulus dissipation along the cell surface (cortex), or the cell barycenter, in which case *η*_(*i,j*)_ = *η*^(*b*)^ is a viscous modulus, representing dissipation within the cell bulk (cytoplasm) (details in the **Supplementary Information**).

First, we conducted simulations for the RUSH-EGFR experiment (**Fig. 1e, f**), considering a down-step in intercellular viscosity on selected clusters of cells (N=4 cells) dispersed in a cell monolayer at equilibrium (**Fig. 4a and Supplementary Information**). We used a set of parameters summarised in **Supplementary Table I**, which were optimised to quantitatively reproduce the experimental results, while remaining within the typical range used to describe MDCK monolayers. In the simulations, a 10-fold decrease (from η=1.4 nN.min.μm^-1^ (30 a.u) to η=0.14 nN.min.μm^-1^ (3 a.u)) in intercellular viscosity leads to a 4-fold increase (from 2.6 ± 0.1 to 10.6 ± 3.7 μm/h; N=100) in the junction elongation velocity (**Fig. 4b, c and Supplementary Video 7**), aligning closely with the experimental values (from 2.6 ± 0.5 to 14.3 ± 1.2 μm/h) (**Fig. 1f**).

Second, we conducted simulations of collective cell migration along the strips (details in **Supplementary Information**). We maintained the same set of parameters while introducing additional conditions: (1) we posited that the inhibition of EGFR by Erlotinib leads to intracellular viscosity equivalent to its downregulation in dense monolayers, given their comparable recruitment levels. We hence set it to η= 1.4 nN.min.μm^-1^ (30 a.u) and (2) we imposed the velocity of the migration front to align with experimental values of 15 μm/h (**Fig. 3g**). Subsequently, we tuned the intercellular viscosity of the Ctrl case to match the experimental results, finding that η= 0.28 nN.min.μm^-1^ (6 a.u) yielded the best quantitative predictions. The **Supplementary Video 7** shows a typical simulation output **(Fig. 4d-f**). Both experimental and simulated data underwent analysis using the same scheme. Notably, a sizeable increase in cell strain (**Fig. 4g**), a decrease in cell velocity (**Fig. 4h**), a reduction in vorticity (**Fig. 4i**), an expansion in spatial correlation length (**Supplementary Fig. 4**), as well as an elevation in cell shape relaxation times t_visc_ (**Fig. 4j, k**) were observed between η= 0.28 nN.min.μm^-1^ (6 a.u) au (Ctrl) and η= 1.4 nN.min.μm^-1^ (30 a.u) (Erlotinib). These quantitative findings closely matched the experimental observations (**Fig. 4l**).

In conclusion, we propose that the E-cadherin-dependent phosphorylation of EGFR fine-tunes the structure of junctional actin, thereby affecting actin dynamics. On a larger scale, it influences junctional viscosity, governing the collective modes of cell migration (**Fig. 2g**). This insight demonstrates that E-cadherin-dependent EGFR activity could regulate the dynamics of collective cell behavior and sheds light on the role of cellular viscous dissipation in collective cell migration, an important aspect that has been understudied.

## Methods

### Cell culture and reagents

MDCK strain II cells were cultured at 37°C with 5% CO_2_ in high-glucose Dulbecco’s Modified Eagle Medium (DMEM, Invitrogen). The medium was supplemented with 10% fetal bovine serum (FBS, Invitrogen), and 100 units/mL of penicillin and 100 μg/mL of streptomycin (Pen-strep, Invitrogen).

To investigate collective migration behaviours, we utilised stable cell lines expressing fluorescent markers or knockout variants of MDCK cells. The following cell lines were employed: wild-type MDCK (MDCK-WT), stably transfected GFP–actin MDCK (MDCK-actin-GFP), stably transfected GFP–E-cadherin MDCK (MDCK-E-cad-GFP) (kindly provided by W. J. Nelson), histone-1–stable GFP MDCK (MDCK-H1-GFP), E-cad KO MDCK (MDCK-Ecad KO) (kindly provided by B. Ladoux, Institut Jacques Monod), and E-cad Rescue MDCK (MDCK-Ecad Res) (kindly provided by P. Kanchanawong, Mechanobiology Institute).

For serum starvation experiments, cells were subjected to serum starvation by incubating them in a growth medium. This medium consisted of high-glucose DMEM lacking FBS, supplemented with 100 units/mL of penicillin and 100 μg/mL of streptomycin. To inhibit EGFR activity, Erlotinib hydrochloride (1μM, Sigma-Aldrich) was employed.

### Plasmids and transfection

The Str-KDEL_SBP-EGFP-EGFR plasmid was generously provided by Dr. David Marc Virshup’s laboratory at Duke NUS. The “SH2-GRB2-tdEOS” plasmid was kindly gifted by Dr. Jay T Groves’ Lab. MDCK-WT cells, with 80% confluence, were transfected with 3 μg of DNA using the Neon electroporation system (Invitrogen), following the manufacturer’s instructions. For Erk activity measurement, EKAREV-NLS expressing MDCK cells were a kind gift from Dr. Tsuyoshi Hirashima’s Laboratory at Mechanobiology Institute, NUS.

### Stamp preparation for collective cell migration on line-patterned strips

Master molds featuring the desired pattern were crafted using SU8-3050 resist on silicon wafers through standard lithography techniques. The pattern employed in this study encompasses a sizable rectangular “reservoir” (approximately 5000 x 700 μm), interconnected with 10 rectangular strips (around 3000 x 400 μm each)^24^. Subsequently, Polydimethylsiloxane (PDMS) stamps were derived from these wafers and utilised for microcontact printing.

The PDMS stamps were incubated with Fibronectin (50μg/ml, Merck) for a duration of 45 minutes, after which they were transferred onto a 35 mm uncoated imaging dish (Ibidi) via microcontact printing. Prior to Fibronectin stamping, the dish had been pre-coated with a layer of PDMS and exposed to UV light for activation. The PDMS stamps were then air-dried within a laminar hood for 10 minutes and delicately pressed against the dish’s bottom for 1 minute. Following microcontact printing, the PDMS stamps were carefully lifted without causing any agitation. The dish bearing the Fibronectin-stamped pattern underwent additional passivation by treating it with a 2% Pluronic F127 solution (Sigma) for 1 hour, aimed at preventing cells from attaching and proliferating in the unstamped areas. Subsequent to passivation, the dishes underwent thorough rinsing with PBS on three times. A PDMS block was strategically positioned atop the microcontact-printed pattern, effectively confining cells within the “reservoir” region. MDCK-H1-GFP, MDCK-Ecad KO, or MDCK-Ecad Res cells were pre-treated with Mitomycin C at a concentration of 10μg/ml (Roche) for a duration of 1 hour to inhibit cell proliferation. These MDCK cells were trypsinised and strategically seeded along the periphery of the PDMS block to cover the “reservoir” area. Once the cells reached confluence on the PDMS block’s sides, the block was gently released, enabling cells to migrate along the strips. Migrating cells were subsequently subjected to specific inhibitors as indicated. The process of live imaging was executed using widefield microscopy (Olympus IX81) with a 10x objective. Throughout imaging, the dishes were maintained within a humidified environment at 37°C with 5% CO_2_. Both phase-contrast and fluorescent images were acquired at intervals of 4 minutes over a duration ranging from 12 to 24 hours.

### Obstacle migration

The design employed for obstacle migration involves a substantial rectangular “reservoir” (approximately 5000 x 700 μm), which is linked to 10 rectangular strips (around 3000 x 400μm each). Each strip features a central circle with a diameter of 200μm. The PDMS stamps crafted from these wafers encompass 200μm diameter circles within each strip. The subsequent preparation steps remain consistent with those outlined in the preceding section. Following contact printing and passivation, the 200μm diameter circles exhibit non-adhesive properties, serving as obstacles during cell migration.

### Single-cell without confinement, single-cell confined to lines, cell trains and cell patches migration

For the migration of single cell lines, wafers with a pattern of 20μm lines were utilised. PDMS stamps were crafted from these wafers and subsequently employed for microcontact printing. The preparation steps mirrored those outlined in the preceding section. Following passivation, the dishes were primed for cell seeding. MDCK-H1-GFP or MDCK-WT cells were trypsinised and their counts were determined prior to seeding. Approximately 4-5 x10^4^ MDCK cells were introduced into the imaging dish and allowed to incubate for 2 hours at 37 °C within a 5% CO_2_ incubator, facilitating full cellular spreading. The migrating cells were then subjected to the indicated inhibitors.

For the migration of cell trains, the employed pattern comprises a large rectangular “reservoir” (approximately 5000 x 700 μm), which is linked to 20 rectangular strips (approximately 3000 x 20μm). The preparation steps mirror those outlined in the preceding section for the creation of line-patterned strips. The migrating cells were subjected to treatment using the specified inhibitors.

In the case of single-cell and cell patch migration, Fibronectin-coated dishes were utilised for direct cell seeding. A quantity of 4-5 x 10^4^ MDCK cells (for single cells) and 3-4 x 10^5^ MDCK cells (for cell patches) were seeded into the imaging dish and incubated at 37 °C within a 5% CO_2_ incubator for 2 hours to allow for complete cell spreading. The migrating cells were subsequently treated with the indicated inhibitors.

Live imaging was conducted using widefield microscopy (Olympus IX81) with either a 10x or 20x objective. The dishes were positioned within a humidified chamber at 37 °C with 5% CO_2_ during the imaging process. Phase-contrast and fluorescent images were captured at 10-minute intervals, spanning a duration ranging from 12 to 24 hours. Migration speeds of individual cells (n > 30) were tracked using either the TrackMate plugin for Image J on phase-contrast images or Imaris for fluorescent nucleus images. The speed of junction deformation at cell-cell junctions was quantified by measuring the lengths of these junctions at each time point.

### Dextran experiments

MDCK-H1-GFP cells (for live imaging) or MDCK-WT cells (for western blotting) were seeded at a low confluence and subjected to overnight serum starvation. Subsequently, 50μg/mL of dextran (00269, 00891, 00894, Sigma-Aldrich) with the specified molecular weight was introduced to the cells before the execution of either western blotting or live imaging. The latter was performed using a widefield microscopy setup (Olympus IX81) equipped with a 20x objective. Cell velocity and confinement ratio were assessed using the Trackmate plugin within Fiji.

### EGFR release experiment

MDCK-WT cells were transfected with the RUSH plasmid Str-KDEL_SBP-EGFP-EGFR. Following transfection, the cells were plated onto an Ibidi imaging dish pre-coated with Fibronectin and placed in a complete medium at 37°C with 5% CO_2_. Once the cells reached confluency, overnight serum starvation was conducted. Subsequently, EGFR-GFP was liberated from the endoplasmic reticulum (ER) through the addition of 40mM biotin. Live imaging was carried out utilising a spinning-disc confocal microscope (Yokogawa CSU-W1) attached to a Nikon Eclipse Ti-E inverted microscope body, equipped with a 60x NA1.3 water lens. Fluorescent images were captured both prior to and subsequent to the biotin introduction, at 5-minute intervals, spanning a duration of 6 hours. The speed of deformation at cell-cell junctions was quantified by measuring the lengths of these junctions at each time point, both 30 minutes before and 30 minutes after the release of EGFR. The measurements were then averaged over this 30-minute period.

### Western blotting

Cells were incubated on ice with RIPA lysis buffer (Sigma), supplemented with protease and phosphatase inhibitor cocktails (Sigma). Subsequently, lysates underwent SDS-PAGE and were transferred onto nitrocellulose membranes. The membranes were then blocked using 5% BSA and 0.05% Tween 20 in TBS, followed by incubation with the specified primary antibodies. Detection of immune complexes was achieved using appropriate HRP-conjugated secondary antibodies (Cell Signaling Technologies) and an enhanced chemiluminescence reagent (Clarity ECL, BioRad). Protein band intensities were quantified using ImageJ Software. The primary antibodies employed were pEGFRY845 (44784G, Thermo Fisher Scientific), pEGFRY1068 (2234, Cell Signaling Technologies), pEGFRY1173 (4407, Cell Signaling Technologies), EGFR (2232, Cell Signaling Technologies), RhoA (sc418, Santa Cruz), Cdc42 (ab187643, Abcam), Rac1 (610651, BD Biosciences) and β-actin (MA515739, Thermo Fisher Scientific).

To assess Rho family GTPases activity, pull-down assay using GST-PBD and GST-RBD were performed on cell lysates as described previously^25^.

### Immunofluorescence

After a collective cell migration period of 12-24 hours, MDCK cells were fixed using pre-warmed 4% paraformaldehyde (PFA) in phosphate-buffered saline (PBS) at 37 °C for 15 minutes. Subsequently, they were permeabilised with 0.2% Triton X-100 in TBS for 30 minutes at room temperature. Samples were then blocked using 1% BSA in TBS for 1 hour. The cells were incubated overnight at 4 °C with primary antibodies: rabbit anti-phospho-EGFR (Y845) polyclonal antibody (44-784G, Thermo Fisher Scientific, diluted 1:200); Purified Mouse anti-E-Cadherin monoclonal antibody (Clone 36) (610181, BD Transduction Laboratories); WAVE2 antibody (H-110) (sc-33548, Santa Cruz); Arp3 antibody (A5979, Sigma-Aldrich); Phospho-Myosin Light Chain 2 (Ser19) antibody (3671, Cell Signaling Technologies, diluted 1:50) according to the manufacturer’s instructions. Following three washes with PBS at 10-minute intervals, cells were incubated with secondary antibodies (anti-mouse Alexa555-conjugated secondary antibody and anti-Rabbit Alexa647-conjugated secondary antibody, both from Thermo Fisher Scientific) along with Alexa405-coupled phalloidin (Invitrogen) in darkness at room temperature for 1 hour. Subsequently, cells were rinsed with PBS and prepared for imaging acquisition. Confocal images were captured in 3D stacks using a spinning-disc confocal microscope (Yokogawa CSU-W1) mounted on a Nikon Eclipse Ti-E inverted microscope body, equipped with a 100x NA1.5 or 60x NA1.3 lens.

### Real-time quantitative PCR (qPCR)

The entire RNA was isolated from a single well of a 6-well plate using the RNeasy Plus Micro Kit (QIAGEN), following the guidelines provided by the manufacturer. A quantity of 450ng of total RNA was employed to generate the cDNA utilising the cDNA synthesis kit (SensiFAST™). For qPCR analysis, the FastStart Universal SYBR Green Master (ROX) mix was employed on a CFX96 Touch Real-Time PCR detection system (Bio-Rad). GAPDH was employed as the internal reference gene.

### Live cell spreading on E-cadherin-coated surface

Disks with a diameter of 25μm were photopatterned onto glass coverslips, following the previously described method^26^. These patterns were then coated overnight at 4°C with a recombinant E-cadherin Fc Tag protein (10204, Sino Biological) at a concentration of 20μg/mL, and subsequently gently washed with PBS. The patterned coverslips were mounted within imaging chambers.

MDCK-WT cells were transfected with the SH2-GRB2-TdEos plasmid, and after selection with 500 μg/mL Geneticin (10131035, Thermo Fisher Scientific), a stable cell line was established following sorting using an SH800S cell sorter (Sony). MDCK-SH2-GRB2-TdEos or MDCK-Ecad-GFP cells were serum-starved overnight, then seeded onto Ecad-Fc patterns, and allowed to spread for a period of 2 hours before initiating TIRF time-lapse imaging. This imaging was carried out using a Motorised TIRF Module (Nikon) integrated with a Nikon Eclipse Ti-E inverted microscope body.

### Fluorescence recovery after photobleaching (FRAP)

Bleaching was performed on the actin of apical cell-cell junctions for FRAP measurements. The cortical actin recovery time (*t*_*half*_) was calculated by fitting the following exponential function to the recovery curves:

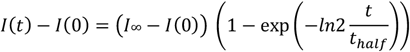

### Laser ablation

An initial image was acquired to determine the precise laser spots. In the case of MDCK-actin-GFP cells, the laser spots were positioned at the apical cell-cell junctions. The pre-acquisition process involved capturing five images at 1-second intervals. During the acquisition phase, a laser power of 60% was utilised, with a duration of approximately 1-2 seconds, targeting the predetermined regions. Subsequently, the post-acquisition stage encompassed the capture of 100 images at 1-second intervals.

To quantify the recoil velocity, the positions of two nodes within the defined junctions were manually tracked using ImageJ software. Following the laser ablation, the temporal evolution of the distance between these two nodes was fitted using a single exponential function. While the double exponential function is commonly employed in other studies, it proved unsuitable for our analysis. In our investigation, the recoil velocity exhibited a gradual nature, and fitting it with a double exponential function yielded unrealistically high speeds. Consequently, we opted to employ a single exponential function for a more appropriate representation.

### Atomic force microscopy (AFM)

AFM experiments were conducted using a Nanowizard IV BioAFM system manufactured by JPK Instruments, Germany. Indentations were performed on randomly chosen cells at the junctional regions. This was achieved using a cantilever (with a nominal k value of 0.03 N/m, provided by Novascan Technologies, Inc., Ames, IA) with an attached polystyrene bead (4.5 μm diameter) at its tip. The applied force was set at 3 nN and a loading rate of 5 μm/s was used.

For each experimental condition, measurements were taken from over 30 cells across three independent trials and subsequently averaged. Young’s modulus values were used to quantitatively describe the cellular stiffness. These values were calculated using the JPK Data Processing Software (JPK Instruments, Germany), which incorporates Hertz’s contact model tailored to spherical indenters (with a diameter of 4.5 μm and a Poisson’s ratio of 0.5). Energy dissipation, representing the heat-based loss of mechanical energy during each indentation cycle by the AFM tip, was determined by assessing the enclosed area between the approach and retraction curves (hysteresis). This phenomenon is largely attributed to frictional and viscous damping within the cell structure at this low speed^27^.

### Erk activity measurement

30000 EKAREV-NLS expressing cells were seeded into a well of culture-inserts 2 wells (81176, ibidi) and allowed to spread for 8 hours. Simultaneously, the insert was then removed and cells were serum starved overnight. Time-lapse FRET images were obtained using a Nikon AX point scanning confocal microscope mounted on a Ti-2 Nikon inverted body. To represent the FRET efficiency, FRET/CFP ratio images were generated after the background was substracted from the original images in the CFP and FRET channel using a matlab code kindly provided by Tsuyoshi Hirashima’s Laboratory at Mechanobiology Institute, NUS. To quantify FRET Ratio for each cells at each timepoints, the Fiji Trackmate plugin was applied to the CFP channel to track each cell position overtime.

### Segmentation

We used Cellpose^28^ for the cell segmentation. We performed an erosion with a 3×3 square kernel to each mask to limit the occurrence of gaps between cells; we disregard objects with areas lower than 20 pixels. We define the cell inertia tensor, with a constant linear weight density along the segmented cell boundaries, i.e. with xx component 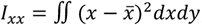, where 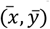 is the position of the cell barycenter. We call *average shape tensor* field the spatially and temporally averaged inertia matrix over all cells within 30×30 pixel-large boxes (corresponding to approx. 10 cells within each), regularly spaced on a spatial grid. The strain field is defined as *ϵ* = log (*λ*_1_/*λ*_2_)/2 where ***λ***_1_ (resp. ***λ***_2_) is the maximum (resp. minimum) eigenvalue of the average shape tensor. For the cell tracking, we used bTrack^29^.

### Simulation of EGFR release experiments

In our EGFR release experiments, cells under investigation are located in the bulk of the tissue, far away from the boundary. In this case, we observe fluctuations in cell edge length but no obvious cell motions, see **Fig. 1e**. To mimic such fluctuations in cell length, we here consider active fluctuations of intercellular tension, 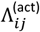, at each cell-cell interface *ij* . These fluctuations contribute to an active force at each vertex,

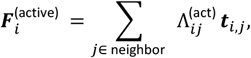

where the summation is over all vertices that connect to the vertex *i*. We assume such tension fluctuations satisfy an Ornstein-Uhlenbeck stochastic dynamic with time correlation^30^,

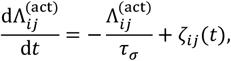

where *τ*_*σ*_is the relaxation time of the active tension and *ζ* _*ij*_ (*t*) are independent Gaussian white noises, satisfying ⟨ *ζ*_*ij*_ (*t*) ⟩ = 0 and 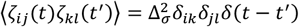 with Δ_*σ*_ being the fluctuation intensity.

We simulated a cell sheet consisting of *N* = 100 cells in a square box of size 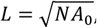, using periodic boundary conditions, see **Supplementary Fig. 1a**. We initialize our simulations from a random Voronoi cell pattern and let the system relax toward a dynamic steady state where the cell elongation parameter and cell motion velocity approach a steady plateau^22^.

To model the effect of the light activation of EGFR and the possibility of a subsequent viscosity modulation, we then randomly selected a small group of four cells in contact and decreased the bulk viscosity 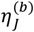of those four cells, from the default value (with 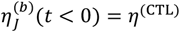) to a lower value (with 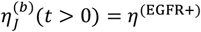) see **Supplementary Fig. 1b**. Further, the viscosity 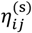 along the cell-cell interface between the vertices *i* and *j* is assumed to be the average viscosity of the two contacting cells (indexed by *J* and *K*) as 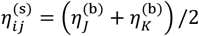.

We compare the junction remodeling velocity, *l*, before and after the drop in viscosity. We find that the ratio of the junction remodeling velocity 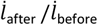 increases with the ratio of the cell viscosity decrease, *η*^(CTL)^/*η*^(EGFR+)^ (**Supplementary Fig. 2**). In particular, the data of *η*^(CTL)^/*η*^(EGFR+)^ = 10 agree with our experiments (**Supplementary Fig. 2**).

We provide the default parameter values for such simulations in **Supplementary Table I**.

### Simulation of collective cell migration experiments

To simulate the collective cell migration experiments, we now turn to collective cells initially confined in a rectangular geometry of size *L*_*x*_ × *L*_*y*_ with 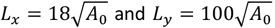 and 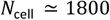, which contains around *N*_cell_ ≃ 1800 cells, see **Supplementary Fig. 3**.

As in the RUSH-EGFR model simulation, we first initialize the simulations using a Voronoi tessellation. We then let the system relax, keeping the bottom boundary fixed and simulating the cell sheet in a confined rectangular geometry to reach a steady state. We next relax the bottom boundary and run the simulations to reach a dynamic steady state.

At the left, top, and right borders, cells are allowed to slip along but adhere to the boundaries; while at the bottom boundary, we imposed vertices to move at a constant speed

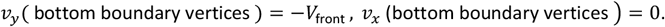

The value of *V*_front_ is fixed at a comparable value to the one measured in experiments. Specifically, we set *V*_front_ = 15 μm/h in simulations.

With the sole migration at the edge (described above), we were not able to recapitulate the formation of vortices similar to the one observed in experiments.

To recapitulate the formation of vortices similar to the one observed in experiments, we turned to a Vicsek-like model of cell motility^23^. Within such model, we associate each cell with an active force 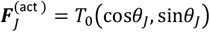 of magnitude *T*_0_ and direction *θ*_*J*_ ; such model mimics the cell motility induced by cell protrusions^23^, with the polar direction *θ*_*J*_ of each cell *J* evolving according to the equation:

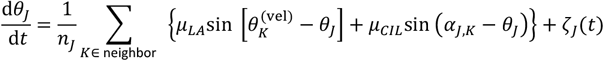

where *μ*_*LA*_ and *μ*_*CIL*_ represent the strengths of local alignment interaction and contact inhibition of locomotion, respectively; 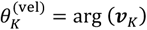 refers to the argument angle of the velocity ***v***_*K*_ of cell *K*; *α*_*J, K*_ = arg (***r***_*J*_ − ***r***_*K*_) denotes the argument angle of the vector pointing from cell *K* to cell *J*; *ζ*_*J*_ (*t*) is a white-noise process with zero mean and variance 2*D*_*r*_. For the cells at the free boundary, we constrain their polar active force direction *θ*_*J*_ to orient normally to the free boundary and toward the free space.

To mimic cell flows from the top boundary (bulk region of MDCK cell sheet), we allow cell divisions on a top region which is within a distance *d* < 5 cell length to the top boundary, see **Supplementary Fig. 3**. We perform cell divisions once cells within such a region exceed an area threshold *A*_div_ = 1.5*A*_0_ = 486*μ*m^2^.

We provide the default parameter values for such simulations in **Supplementary Table II**.

We are interested in and examine the collective cell dynamics in a region near the moving front (within a distance of ∼ 30 cell length to the moving front). Note that to reduce the artificial effect of the top boundary condition on collective cell migration dynamics in a region near the moving front, we have set a sufficiently large scale of the simulated cell monolayer in the vertical direction, i.e., *L*_*y*_ ∼ 100 cell length.

### Data display and statistics

Prism (GraphPad Software) and Matlab (Math Works) were used for data analysis and graph plotting. Graphs were mounted using Adobe Illustrator. ANOVA test and paired or unpaired Student’s t-test were carried out to analyse the significant difference levels.

## Acknowledgements

V.V and MS acknowledge support from MOE grant MOE2016-T3-1-002, NRF grant NRFI2018-07 and Seed funding from MBI

J.-F. R. is hosted at the Laboratoire Adhésion Inflammation (LAI). The project leading to this publication has received funding from France 2030, the French Government program managed by the French National Research Agency (ANR-16-CONV-0001) and from Excellence Initiative of Aix-Marseille University - A*MIDEX. J.-F. R. is also funded by ANR-20-CE30-0023 COVFEFE

## Author contributions

Fu Chaoyu performed the migration experiments, their quantification and wrote the manuscript. F. Dilasser performed the single cell experiments and the dextran experiment, the pull-down assay. Zhao-zhen Lin performed all simulations. Marx Karnat did all the segmentation and tracking, Sound Wai Phow and Tsuyoshi Hirashima helped with the ERK experiments. Hui Ting Ong helped with image analysis. Nai Mui Hoon Brenda performed the AFM experiments. Harini performed the QPCR, Aditya Arora contributed to the understanding of the experimental results. Michael Sheetz initiated the work on EGFR and supervised C.F. Jean-Francois Rupprecht supervised the simulation work and contributed to the manuscript. Sham Tlili performed the relaxation time analysis and contributed to the theoretical part of the work. Virgile Viasnoff designed the experiments, contributed to the manuscript and supervised the work.

## Conflict of Interest

The authors declare no conflict of interest.

## Extended Data Figures

**Extended Data Fig. 1:**
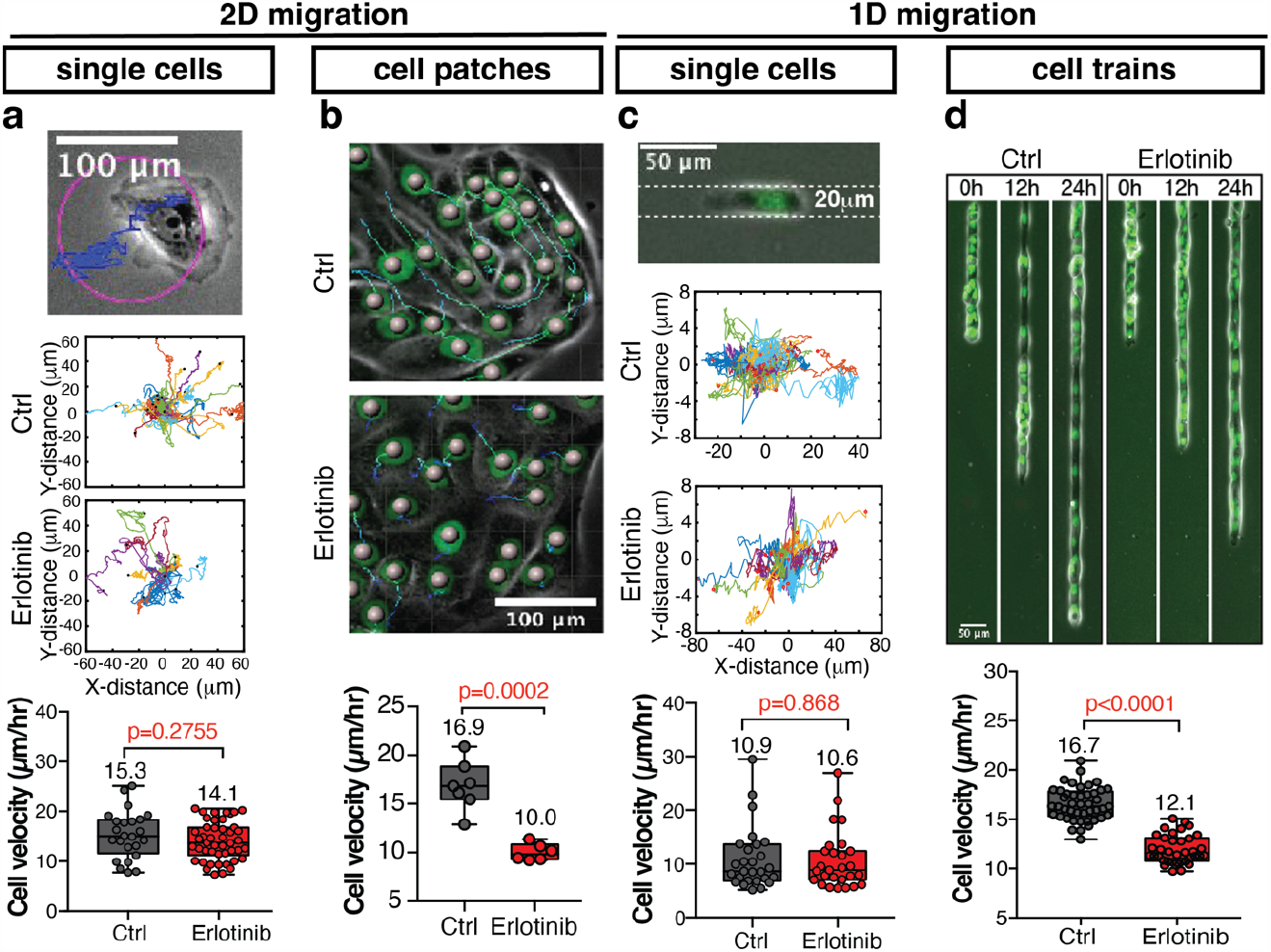
Dephosphorylation of EGFR reduces the dynamics of cell junction deformation without directly altering cell-substrate interactions. **a**.Inhibition of EGFR activity does not disrupt single-cell motility in 2D migration. Representative examples of single-cell 2D migration on fibronectin-coated surfaces (top), Scale Bar: 100 μm. Thirty to forty representative nuclear tracks over 12h in cells randomly migrating under Ctrl and pEGFR inhibition conditions (middle). Average speed of single-cell motility from nuclear movement tracks over 12h under Ctrl and pEGFR inhibition conditions (bottom), n_ctrl_ = 24 cells and n_Erlotinib_ = 46 cells from 3 independent experiments, two-tailed unpaired t-test, p=0.2755. **b**.Representative examples of migration of 2D epithelial patches on fibronectin-coated surfaces under Ctrl and pEGFR inhibition conditions (top), Scale Bar: 100 μm. Average speed of cells in patch migration for 12h from nuclear movement tracks under Ctrl and pEGFR inhibition conditions, n_Ctrl_ = 7 cell patches and n_Erlotinib_ = 6 cell patches from 4 different experiments, two-tailed unpaired t-test, p=0.0002. **c**.Inhibition of EGFR activity does not disturb single-cell motility in 1D migration. Representative examples of single-cell migration on 20 μm fibronectin-coated lines (top), Scale Bar: 50 μm. Thirty representative nuclear tracks over 12h in cells migrating on 20 μm line patterns under Ctrl and pEGFR inhibition conditions (middle). Average speed of single-cell motility on 20 μm line patterns from nuclear movement tracks under Ctrl and pEGFR inhibition conditions (bottom), n_Ctrl_ = 27 cells and n_Erlotinib_ = 27 cells from 3 different experiments, two-tailed unpaired t-test, p=0.868. **d**.Representative examples of migration of 1D epithelial trains on 20 μm fibronectin-coated lines under Ctrl and pEGFR inhibition conditions (top), Scale Bar: 50 μm. Average speed of cells in train migration for 24h on 20 μm line patterns from nuclear movement tracks under Ctrl and pEGFR inhibition conditions, n_Ctrl_ = 44 cell trains and n_Erlotinib_ = 36 cell trains from 3 different experiments, two-tailed unpaired t-test, p<0.0001.

**Extended Data Fig. 2:**
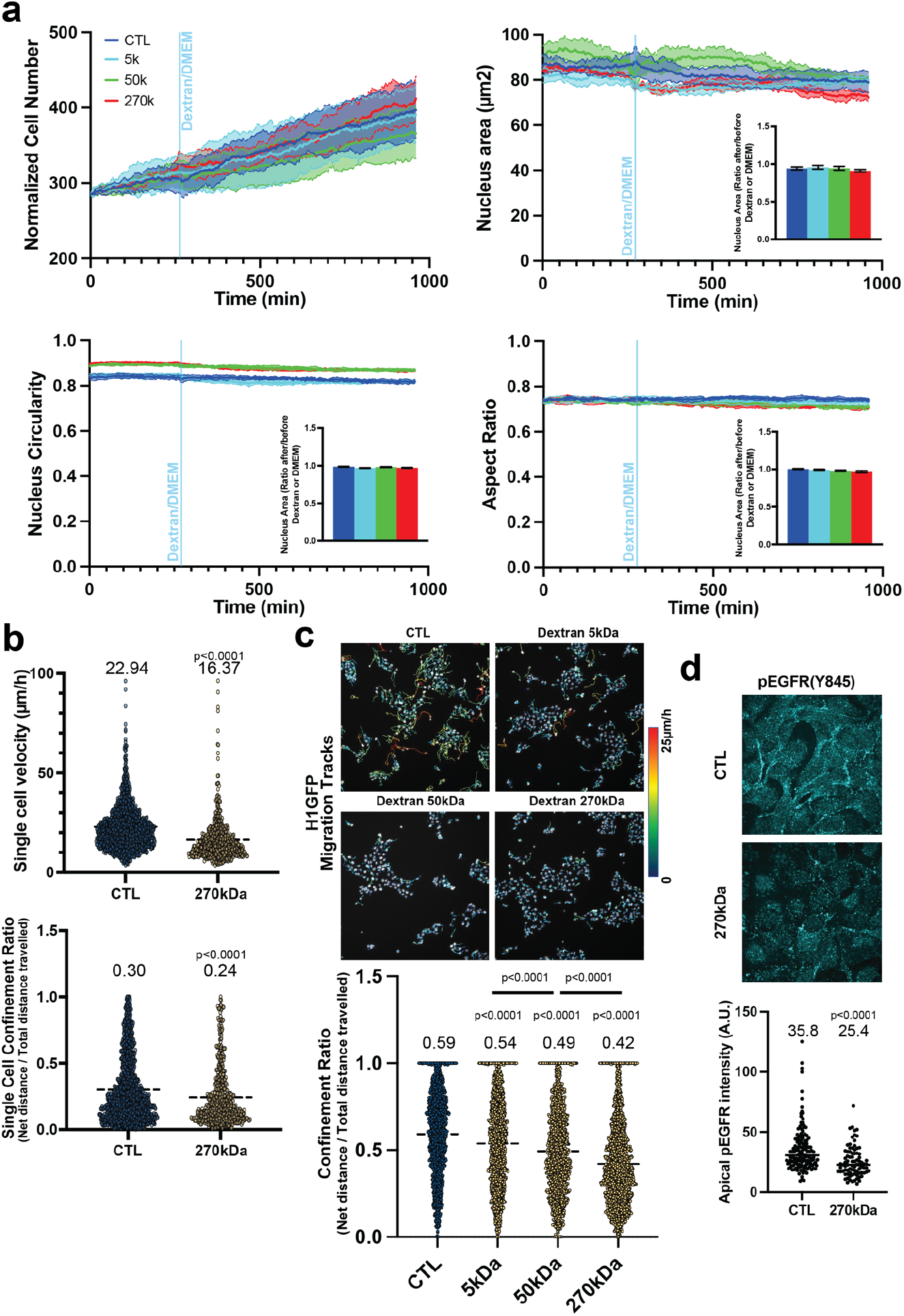
Dextran impact on single and collective cell migration speed and mode without evident signs of osmotic shock. **a**.Quantification of cell proliferation (upper left), nucleus area (upper right), circularity (lower left) and aspect ratio (lower right) on cell patches before and after the addition of dextran of the indicated molecular weight. The light blue line indicates the moment of dextran addition. Four independent experiments yielded consistent results. **b**.Quantification of single-cells velocity and persistence before and after the addition of 270kDa dextran, n_Ctrl_ = 1217 cells and n_270kDa_ = 543 cells from 3 different experiments, two-tailed unpaired t-test, p<0.0001. **c**.Representative images of cell tracking with or without the presence of dextran of the indicated molecular weight (up). Quantification of individual cell persistence within cell patches under control or indicated molecular weight dextran conditions (down), n_Ctrl_ = 1762 cells, n_5kDa_ = 1770 cells, n_50kDa_ = 1424 cells and n_270kDa_ = 1968 cells from 3 different experiments, Ordinary one-way ANOVA Tukey’s test, p<0.0001. **d**.Confocal images (up) and quantification (down) of pEGFR(Y845) junctional fluorescence intensities under control and 270kDa dextran conditions, n_Ctrl_ = 143 cell junctions and n_270kDa_ = 84 cell junctions from 3 different experiments, two-tailed unpaired t-test, p<0.0001.

**Extended Data Fig. 3:**
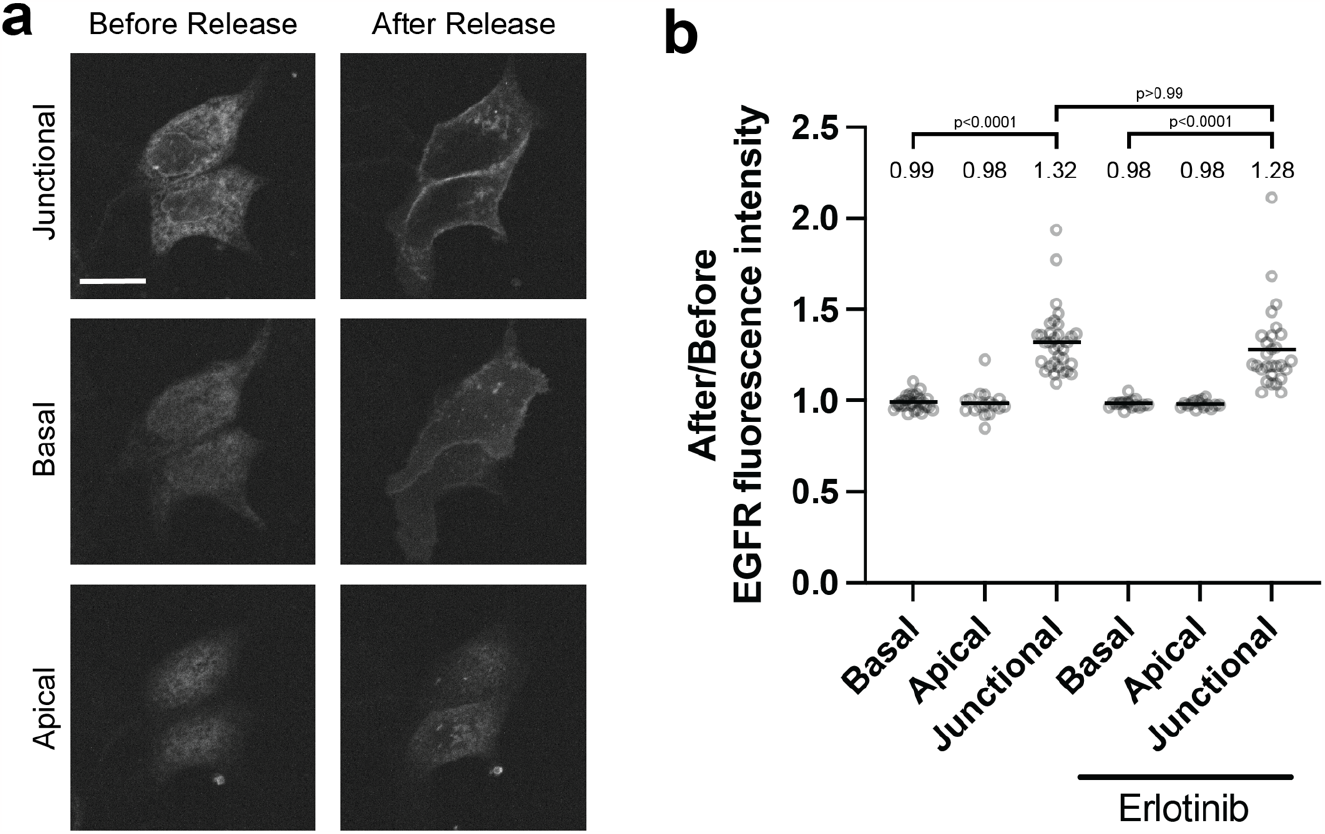
EGFR preferentially translocates to cell-cell junction after release from the Endoplasmic Reticulum (ER). **a-b**. Confocal images (**a**) and corresponding quantification (**b**) of EGFR-GFP fluorescence intensities at the junctional, basal and apical side of the cells, both before and after release from the endoplasmic reticulum. Scale Bar: 20 μm. n_Ctrl Basal_ = 27 cells, n_Ctrl Apical_ = 17 cells, n_Ctrl Junctional_ = 33 cell junctions, n_Erlotinib Basal_ = 17 cells, n_Erlotinib Apical_ = 15 cells and n_Erlotinib Junctional_ = 28 cell junctions from 3 different experiments, Ordinary one-way ANOVA Tukey’s test.

**Extended Data Fig. 4:**
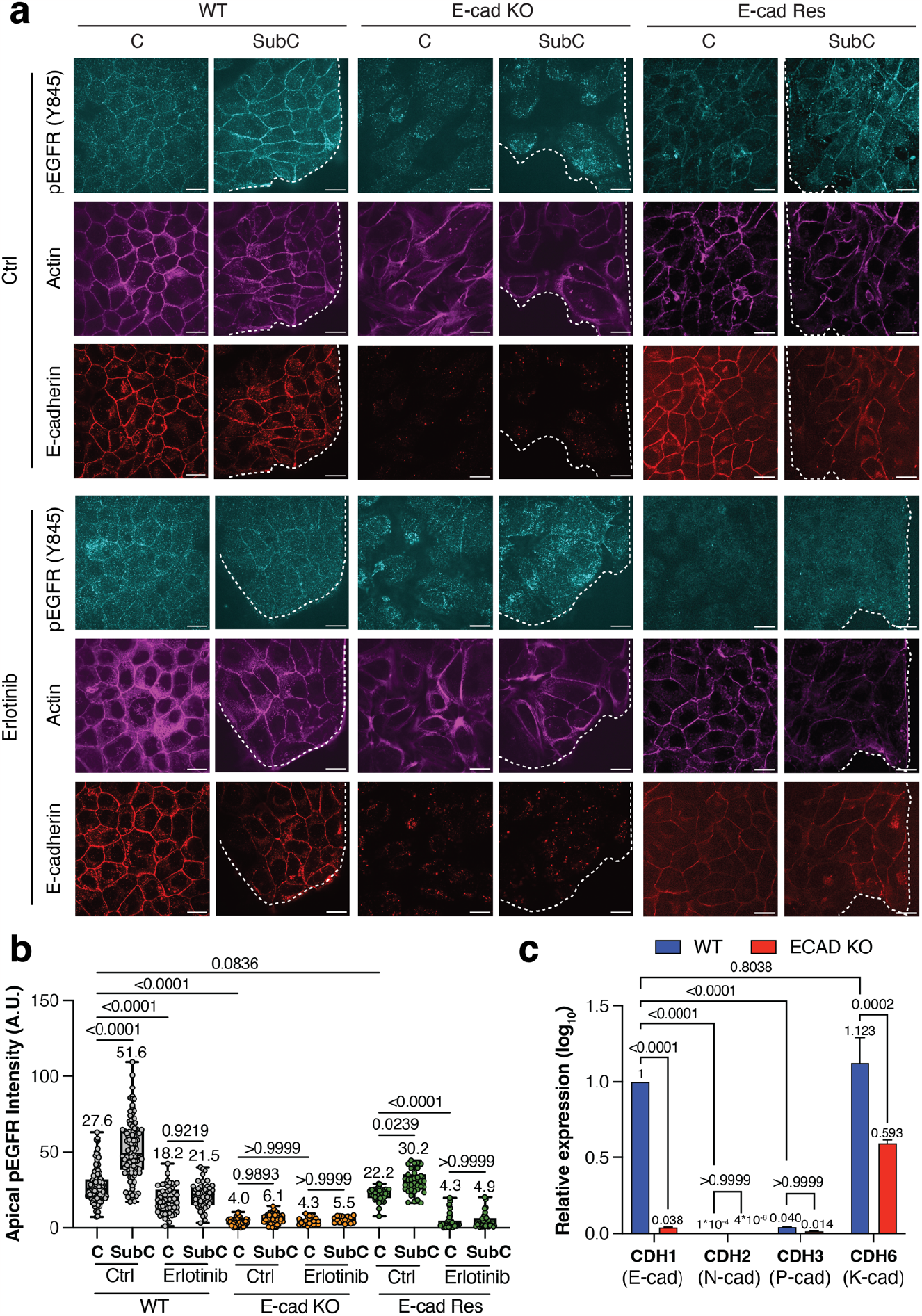
E-cadherin junctions govern phosphorylation of EGFR (Y845) at the apical side of MDCK tissues. **a**.Immunostaining of pEGFR (Y845), actin and E-cadherin in wild-type (WT), E-cadherin knock-out (Ecad-KO) and Ecad-KO-rescued (Ecad-Res) MDCK cells on the apical side of confluent (C, left) and sub-confluent (SubC, right) regions under control and pEGFR inhibition (Erlotinib) conditions. The white dotted line indicates the leading front of the patches. Scalr Bar: 20 μm. **b**.Quantification of apical pEGFR in WT, Ecad-KO and Ecad-Res MDCK cells. For WT: n_Ctrl, C_ = 122 cell junctions, n_Ctrl, SubC_ = 106 cell junctions from 4 different experiments, n_Erlotinib, C_ = 51 cell junctions, n_Erlotinib, SubC_ = 44 cell junctions from 3 different experiments; For Ecad-KO: n_Ctrl, C_ = 67 cell junctions, n_Ctrl, SubC_ = 61 cell junctions from 3 different experiments, n_Erlotinib, C_ = 24 cell junctions, n_Erlotinib, SubC_ = 16 cell junctions from 3 different experiments; For Ecad-Res: n_Ctrl, C_ = 47 cell junctions, n_Ctrl, SubC_ = 34 cell junctions from 3 different experiments, n_Erlotinib, C_ = 41 cell junctions, n_Erlotinib, SubC_ = 32 cell junctions from 3 different experiments, Ordinary one-way ANOVA Tukey’s test. **c**.RT-qPCR results showing the relative expression of CDH1 (E-cad), CDH2 (N-cad), CDH3 (P-cad) and CDH6 (K-cad) in WT and Ecad-KO MDCK cells from 3 different experiments.

**Extended Data Fig. 5:**
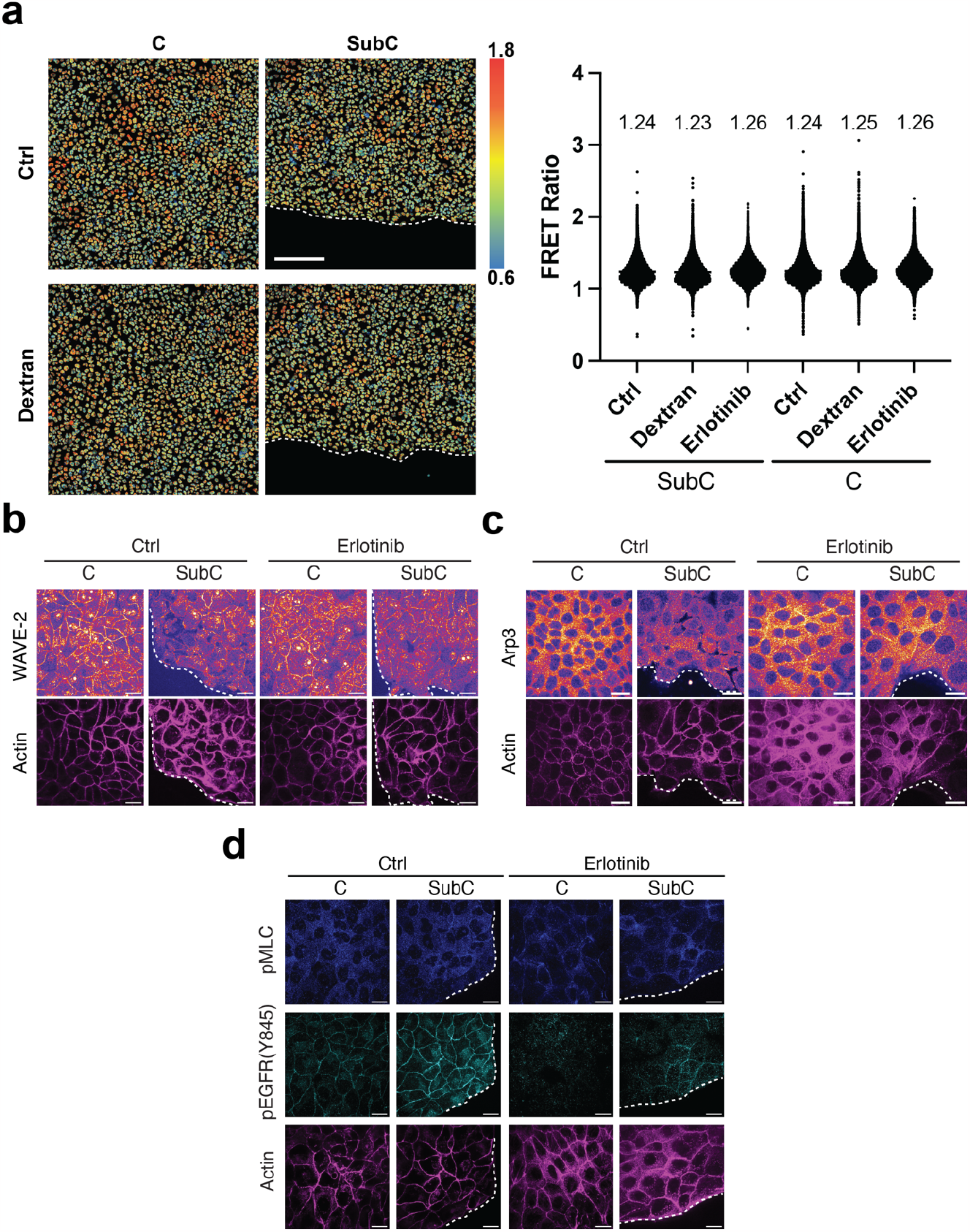
Molecular components underlying EGFR phosphorylation in response to cell junction deformation. **a**. Representative images and corresponding quantification of FRET Ratio in confluent (C, left) and sub-confluent (SubC, right) regions of migrating MDCK monolayers, with or without EGFR inhibition (Erlotinib), n_Ctrl, SubC_ = 27014 cells, n_Dextran, SubC_ = 27766 cells, n_Erlotinib, SubC_ = 15316 cells, n_Ctrl, C_ = 43852 cells, n_Dextran, C_ = 43079 cells and n_Erlotinib, C_ = 27305 cells from 3 independent experiments. Scale Bar: 200 μm. **b-c**. Immunostaining of WAVE-2 and actin (**b**), Arp3 and actin (**c**) on the apical side of confluent (C, left) and sub-confluent (SubC, right) regions under Control and pEGFR inhibition (Erlotinib) conditions. Scale Bar: 20 μm. Two independent experiments yielded consistent results. **d**. Immunostaining of pMLC, pEGFR (Y845) and actin on the apical side of confluent (C, left) and sub-confluent (SubC, right) regions under control and pEGFR inhibition (Erlotinib) conditions. Scale Bar: 20 μm. Two independent experiments yielded consistent results.

**Extended Data Fig. 6:**
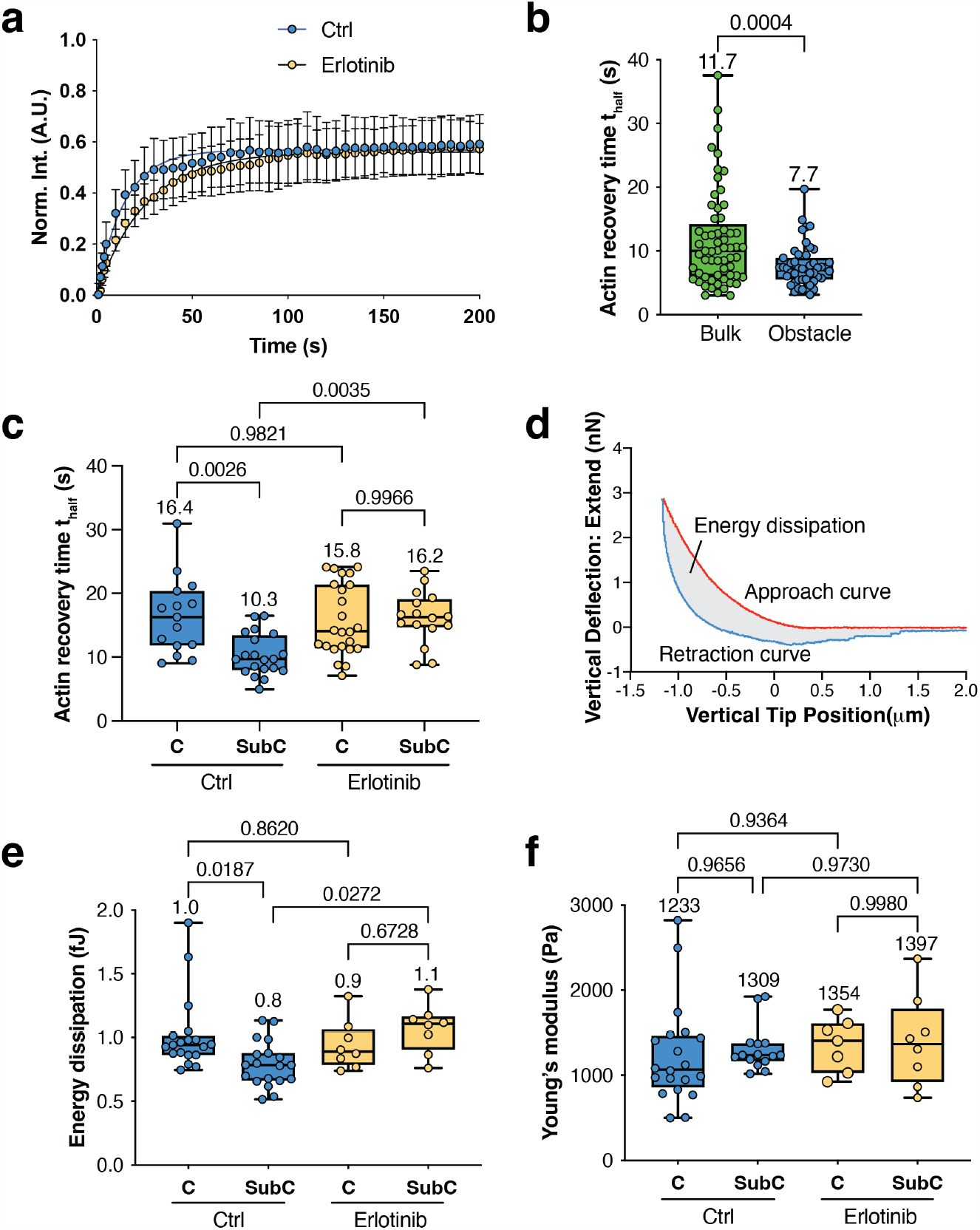
Regulation of actin dynamics and junctional viscoelastic properties by EGFR phosphorylation without impact on junctional tension. **a**.Fluorescence Recovery After Photobleaching (FRAP) experiments of actin were conducted at the apical regions of cell-cell junctions under both control and pEGFR inhibition (Erlotinib) conditions. The data were normalized and fitted with best-fit curves. **b**.Actin dynamics were assessed in cells encircling obstacles and in bulk regions. n_Bulk_ = 61 cell junctions and n_Obstacle_ = 47 cell junctions from 3 independent experiments, two-tailed unpaired t-test, p=0.0004. **c**.Measurements of actin dynamics in migrating MDCK monolayers were performed in both confluent (C, left) and sub-confluent (SubC, right) regions under control and pEGFR inhibition (Erlotinib) conditions. n_Ctrl, C_ = 15 cell junctions, n_Ctrl, SubC_ = 20 cell junctions from 2 independent experiments, n_Erlotinib, C_ = 24 cell junctions, n_Erlotinib, SubC_ = 16 cell junctions from 3 independent experiments, Ordinary one-way ANOVA Tukey’s test. **d**.Typical force-displacement curves from Atomic Force Microscopy (AFM) force measurements of MDCK monolayers. The area enclosed between the approach and retraction curves, corresponds to energy dissipation. **e-f**. Measurements of energy dissipation (**e**) and Young’s modulus (**f**) of migrating MDCK monolayers in confluent (C, left) and sub-confluent (SubC, right) regions under control and pEGFR inhibition (Erlotinib) conditions. For energy dissipation: n_Ctrl, C_ = 19 cell junctions, n_Ctrl, SubC_ = 19 cell junctions from 3 independent experiments, n_Erlotinib, C_ = 8 cell junctions, n_Erlotinib, SubC_ = 8 cell junctions from 3 independent experiments, Ordinary one-way ANOVA Tukey’s test. For Young’s modulus: n_Ctrl, C_ = 20 cell junctions, n_Ctrl, SubC_ = 15 cell junctions from 3 independent experiments, n_Erlotinib, C_ = 7 cell junctions, n_Erlotinib, SubC_ = 8 cell junctions from 3 independent experiments, Ordinary one-way ANOVA Tukey’s test.

## Supplementary Discussion

### E-Cadherin - EGFR - Rac1 - Wave2 - Arp2/3 interplay and relevance at cell-cell junctions

In this study, we elucidated a molecular signaling pathway that plays a crucial role in regulating the viscosity of intercellular junctions. Our findings demonstrate that the trans binding of E-cadherin within highly dynamic cell-cell junctions triggers the activation of EGFR and Rac1, aligning with previous research^7, 31–33^. Rac1 has been well-established to play a major role in Wave2 complex recruitment and activation^34^. Specifically, E-cadherins activate Rac1 at the cell-cell junction, contributing to junction stabilization through Wave2^35, 36^.

Contrary to existing studies conducted on highly confluent monolayers characterized by mature contacts and non-dynamic cell-cell junctions, our investigation focuses on sub-confluent patches exhibiting elevated apical pEGFR and Rac1-GTP levels. In our unique context of high dynamics, we proposed that the signaling pathways and molecular actors may differ. Notably, Rac1 activation, which has been linked to E-cadherin adhesion disruptions^37^, could potentially contribute to the rapid turnover of E-cadherin adhesions in highly dynamic cell-cell junctions, thereby hindering Wave2 recruitment for junction stabilization.

Alternatively, we hypothesize that the impact of pEGFR at the junction is non-local, coupling with recruitment at the basal part of cells and creating competition with the junctional pool. Unfortunately, due to limitations in the available imaging techniques, we were unable to discern the cortex bound to the cytoplasmic fraction. Consequently, while our paper establishes a correlation between the recruitment levels of various proteins, we refrain from asserting a direct causal link.

### Captions of the Supplementary videos

**Supplementary Video 1:**

Cell-cell reorganization in representative patches of MDCK-H1-GFP cells under control and pEGFR-inhibited (Erlotinib at 1μM) conditions recorded at 12 frames/hour. Scale Bar: 100μm.

**Supplementary Video 2:**

Cell patches migration of MDCK-H1-GFP cells with or without dextran addition recorded at 12 frames/hour. Scale Bar: 100μm.

**Supplementary Video 3:**

MDCK-WT monolayers with mosaic expression of RUSH-EGFR-GFP. Junction elongation before and after addition of biotin and subsequent release of EGFR from the endoplasmic reticulum under control and pEGFR-inhibited (Erlotinib at 1μM) conditions recorded at 12 frames/hour. Scale Bar: 20μm.

**Supplementary Video 4:**

Cells encircling obstacles and in bulk regions under control and pEGFR-inhibited (Erlotinib at 1μM) conditions recorded at 15 frames/hour. Scale Bar: 50μm.

**Supplementary Video 5:**

Cells migrating on 400 μm width line strips under control and pEGFR-inhibited (Erlotinib at 1μM) conditions recorded at 15 frames/hour. Scale Bar: 100μm.

**Supplementary Video 6:**

The vorticity of the collective flow for the migrating cells under control and pEGFR-inhibited (Erlotinib at 1μM) conditions recorded at 15 frames/hour.

**Supplementary Video 7:**

Vertex-model simulations under control (η=6) and pEGFR-inhibited (Erlotinib at 1μM) (η=30) conditions. Other parameters see Supplementary Table II.

## Notes

### Competing Interest Statement

The authors have declared no competing interest.

## Reference

1. Martin, A.C., Kaschube, M. & Wieschaus, E.F. Pulsed contractions of an actin-myosin network drive apical constriction. Nature 457, 495–499 (2009).

2. Bertet, C., Sulak, L. & Lecuit, T. Myosin-dependent junction remodelling controls planar cell intercalation and axis elongation. Nature 429, 667–671 (2004).

3. Reffay, M. et al. Interplay of RhoA and mechanical forces in collective cell migration driven by leader cells. Nat Cell Biol 16, 217–223 (2014).

4. Ridley, A.J. Rho GTPase signalling in cell migration. Curr Opin Cell Biol 36, 103–112 (2015).

5. Mason, F.M., Xie, S., Vasquez, C.G., Tworoger, M. & Martin, A.C. RhoA GTPase inhibition organizes contraction during epithelial morphogenesis. J Cell Biol 214, 603–617 (2016).

6. Heer, N.C. & Martin, A.C. Tension, contraction and tissue morphogenesis. Development 144, 4249–4260 (2017).

7. Pece, S. & Gutkind, J.S. Signaling from E-cadherins to the MAPK pathway by the recruitment and activation of epidermal growth factor receptors upon cell-cell contact formation. J Biol Chem 275, 41227–41233 (2000).

8. Fu, C., Arora, A., Engl, W., Sheetz, M. & Viasnoff, V. Cooperative regulation of adherens junction expansion through epidermal growth factor receptor activation. J Cell Sci 135 (2022).

9. Xu, K.P., Yin, J. & Yu, F.S. SRC-family tyrosine kinases in wound- and ligand-induced epidermal growth factor receptor activation in human corneal epithelial cells. Invest Ophthalmol Vis Sci 47, 2832–2839 (2006).

10. Boncompain, G. et al. Synchronization of secretory protein traffic in populations of cells. Nat Methods 9, 493–498 (2012).

11. Saxena, M. et al. EGFR and HER2 activate rigidity sensing only on rigid matrices. Nat Mater 16, 775–781 (2017).

12. Antczak, C., Mahida, J.P., Bhinder, B., Calder, P.A. & Djaballah, H. A high-content biosensor-based screen identifies cell-permeable activators and inhibitors of EGFR function: implications in drug discovery. J Biomol Screen 17, 885–899 (2012).

13. Hino, N. et al. ERK-Mediated Mechanochemical Waves Direct Collective Cell Polarization. Dev Cell 53, 646–660 e648 (2020).

14. Hall, A. Rho GTPases and the actin cytoskeleton. Science 279, 509–514 (1998).

15. Verma, S. et al. A WAVE2-Arp2/3 actin nucleator apparatus supports junctional tension at the epithelial zonula adherens. Mol Biol Cell 23, 4601–4610 (2012).

16. Arora, A. et al. Cortical ductility governs cell-cell adhesion mechanics. bioRxiv preprint (2023).

17. Tlili, S. et al. Migrating Epithelial Monolayer Flows Like a Maxwell Viscoelastic Liquid. Phys Rev Lett 125, 088102 (2020).

18. Honda, H. & Eguchi, G. How much does the cell boundary contract in a monolayered cell sheet? J Theor Biol 84, 575–588 (1980).

19. Barton, D.L., Henkes, S., Weijer, C.J. & Sknepnek, R. Active Vertex Model for cell-resolution description of epithelial tissue mechanics. PLoS Comput Biol 13, e1005569 (2017).

20. Fletcher, A.G., Osterfield, M., Baker, R.E. & Shvartsman, S.Y. Vertex models of epithelial morphogenesis. Biophys J 106, 2291–2304 (2014).

21. Alt, S., Ganguly, P. & Salbreux, G. Vertex models: from cell mechanics to tissue morphogenesis. Philos Trans R Soc Lond B Biol Sci 372 (2017).

22. Lin, S.Z., Merkel, M. & Rupprecht, J.F. Structure and Rheology in Vertex Models under Cell-Shape-Dependent Active Stresses. Phys Rev Lett 130, 058202 (2023).

23. Lin, S.Z., Ye, S., Xu, G.K., Li, B. & Feng, X.Q. Dynamic Migration Modes of Collective Cells. Biophys J 115, 1826–1835 (2018).

24. Vedula, S.R. et al. Emerging modes of collective cell migration induced by geometrical constraints. Proc Natl Acad Sci U S A 109, 12974–12979 (2012).

25. Guilluy, C., Dubash, A.D. & Garcia-Mata, R. Analysis of RhoA and Rho GEF activity in whole cells and the cell nucleus. Nat Protoc 6, 2050–2060 (2011).

26. Azioune, A., Storch, M., Bornens, M., Thery, M. & Piel, M. Simple and rapid process for single cell micro-patterning. Lab Chip 9, 1640–1642 (2009).

27. Yang, R. et al. Characterization of mechanical behavior of an epithelial monolayer in response to epidermal growth factor stimulation. Exp Cell Res 318, 521–526 (2012).

28. Stringer, C., Wang, T., Michaelos, M. & Pachitariu, M. Cellpose: a generalist algorithm for cellular segmentation. Nat Methods 18, 100–106 (2021).

29. Ulicna, K., Vallardi, G., Charras, G. & Lowe, A.R. Automated Deep Lineage Tree Analysis Using a Bayesian Single Cell Tracking Approach. Frontiers in Computer Science 3 (2021).

30. Tlili, S. et al. Shaping the zebrafish myotome by intertissue friction and active stress. Proc Natl Acad Sci U S A 116, 25430–25439 (2019).

31. Nakagawa, M., Fukata, M., Yamaga, M., Itoh, N. & Kaibuchi, K. Recruitment and activation of Rac1 by the formation of E-cadherin-mediated cell-cell adhesion sites. J Cell Sci 114, 1829–1838 (2001).

32. Kovacs, E.M., Ali, R.G., McCormack, A.J. & Yap, A.S. E-cadherin homophilic ligation directly signals through Rac and phosphatidylinositol 3-kinase to regulate adhesive contacts. J Biol Chem 277, 6708–6718 (2002).

33. Betson, M., Lozano, E., Zhang, J. & Braga, V.M. Rac activation upon cell-cell contact formation is dependent on signaling from the epidermal growth factor receptor. J Biol Chem 277, 36962–36969 (2002).

34. Rottner, K., Faix, J., Bogdan, S., Linder, S. & Kerkhoff, E. Actin assembly mechanisms at a glance. J Cell Sci 130, 3427–3435 (2017).

35. Yamazaki, D., Oikawa, T. & Takenawa, T. Rac-WAVE-mediated actin reorganization is required for organization and maintenance of cell-cell adhesion. J Cell Sci 120, 86–100 (2007).

36. Erasmus, J.C., Welsh, N.J. & Braga, V.M. Cooperation of distinct Rac-dependent pathways to stabilise E-cadherin adhesion. Cell Signal 27, 1905–1913 (2015).

37. Lozano, E., Frasa, M.A., Smolarczyk, K., Knaus, U.G. & Braga, V.M. PAK is required for the disruption of E-cadherin adhesion by the small GTPase Rac. J Cell Sci 121, 933–938 (2008).

