## Supplementary info for "E-cadherin-dependent phosphorylation of EGFR governs a homeostatic feedback loop controlling intercellular junction viscosity and collective migration modes"

(Dated: November 23, 2023)

**Contents**


**I. Model Methods** 2
A. Vertex model description 2
B. Viscous forces 2
C. Numerical scheme 4
D. Evaluation of tissue viscoelastic relaxation time 9
1. Tensor maps 9
2. Extraction of a viscoelastic relaxation time 11

**References** 13

### I. MODEL METHODS

#### A. Vertex model description

We use a cell-based computational model, called vertex model [1–3]. Vertex models describe epithelial tissues as polygonal tilings, see Supplementary Figure 1a; the dynamics of the polygon vertices correspond to a force balance equation.

Here, we consider a force balance equation in the form:

$$\underbrace{\boldsymbol{F}_i^{(\text{friction})}}_{\text{friction}} + \underbrace{\boldsymbol{F}_i^{(\text{viscous})}}_{\text{viscosity}} + \underbrace{\boldsymbol{F}_i^{(\text{elastic})}}_{\text{elasticity}} + \underbrace{\boldsymbol{F}_i^{(\text{active})}}_{\text{activity}} = \boldsymbol{0}, \tag{S1}$$

which accounts for a balance between:

1. friction forces between the monolayer and the substrate (typically mediated through adhesion complexes), expressed as:

$$\boldsymbol{F}_i^{(\text{friction})} = -\xi \boldsymbol{v}_i, \tag{S2}$$

   with $\xi$ being the friction coefficient and $\boldsymbol{v}_i = \mathrm{d}\boldsymbol{r}_i/\mathrm{d}t$ being the velocity of vertex $i$.

2. the cellular viscous forces, stemming from intercellular junctional viscosity and intracellular cytoplasm viscosity. The expression of $\boldsymbol{F}_i^{(\text{viscous})}$ is derived in the Method section, I B.

3. elastic forces corresponding to mechanical regulation of the cell shape, which, following [2, 4, 5], we consider to derive from a mechanical energy $E$ such that $\boldsymbol{F}_i^{(\text{elastic})} = -\partial E/\partial \boldsymbol{r}_i$. We express the mechanical energy of an epithelial cell sheet as [2, 4–6],

$$E = \underbrace{\sum_J \frac{1}{2} K_A (A_J - A_0)^2}_{\text{area elasticity}} + \underbrace{\sum_J \frac{1}{2} K_P (P_J - P_0)^2}_{\text{perimeter elasticity}}, \tag{S3}$$

   where the two terms account for the cell area elasticity and the cell perimeter elasticity, respectively. In detail, $K_A$ and $K_P$ are the area stiffness and perimeter stiffness of cells, respectively; $A_J$ and $P_J$ are the area and the perimeter of the $J-$th cell, respectively; $A_0$ and $P_0$ are the preferred area and the preferred perimeter, respectively.

4. active forces due to fluctuations in intercellular junction tensions or due to cell-substrate tractions (see main text, Method section).

#### B. Viscous forces

We consider two types of cellular viscosity, i.e., cell–cell interfacial viscosity ($\eta_{ij}^{(\text{s})}$) and cell bulk viscosity ($\eta_J^{(\text{b})}$), as sketched in Supplementary Figure 1. The viscosity $\eta_{ij}^{(\text{s})}$ along the cell–cell interface between the vertices $i$ and $j$ is assumed to be the average viscosity of the two contacting cells (indexed by $J$ and $K$) as $\eta_{ij}^{(\text{s})} = (\eta_J^{(\text{b})} + \eta_K^{(\text{b})})/2$. The resulting viscous force is expressed as

$$\boldsymbol{F}_i^{(\text{viscous})} = \sum_{j \in V_i} \Lambda_{ij}^{(\text{viscous})} \boldsymbol{t}_{i,j} + \sum_{J \in C_i} \Lambda_{iJ}^{(\text{viscous})} \boldsymbol{t}_{i,J}, \tag{S4}$$

The first term in Eq. (S4) accounts for the viscosity of intercellular junctions (Supplementary Figure 1b), and the summation $\sum_{j \in V_i}$ is over all neighboring vertices $V_i$ of vertex $i$. We express the viscous tension $\Lambda_{ij}^{(\text{viscous})}$ of the cell—cell interface $ij$ as,

$$\Lambda_{ij}^{(\text{viscous})} = \eta_{ij}^{(\text{s})} \dot{l}_{ij}, \tag{S5}$$

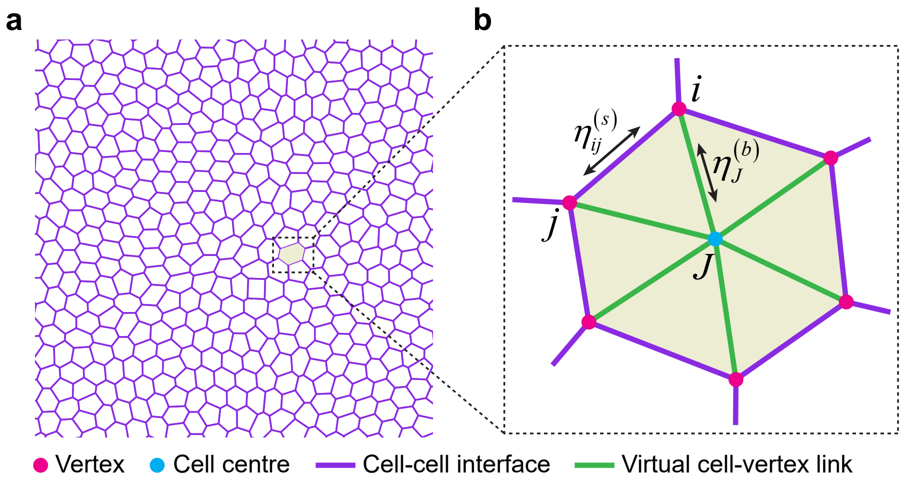


Supplementary Figure 1: Sketch of the cell viscosity implemented in our vertex model. (a) Geometric description of an epithelial cell sheet by a two-dimensional tiling of polygons. (b) In our model, we consider two kinds of cell viscosity, including cell–cell interfacial viscosity, $\eta_{ij}$, and cell bulk viscosity, $\eta_J$.

where $l_{ij} = |\boldsymbol{r}_i - \boldsymbol{r}_j|$ is the length of the cell–cell interface $ij$; $\dot{l}_{ij} = \mathrm{d}l_{ij}/\mathrm{d}t$ is the elongation rate of the cell–cell interface $ij$; $\eta^{(\mathrm{s})}_{ij}$ is the viscous coefficient of cell–cell interface $ij$. Since $\dot{l}_{ij} = \boldsymbol{t}_{i,j} \cdot (\boldsymbol{v}_j - \boldsymbol{v}_i)$ with $\boldsymbol{t}_{i,j} = (\boldsymbol{r}_j - \boldsymbol{r}_i)/l_{ij}$ being a unit vector along the cell–cell interface $ij$, the viscous tension $\Lambda^{(\mathrm{viscous})}_{ij}$ can be further expressed as,

$$\Lambda^{(\mathrm{viscous})}_{ij} = \eta^{(\mathrm{s})}_{ij} \boldsymbol{t}_{i,j} \cdot (\boldsymbol{v}_j - \boldsymbol{v}_i)\,. \tag{S6}$$

The second term in Eq. (S4) accounts for the viscosity of cell cytoplasm (Supplementary Figure 1b), and the summation $\sum_{J \in C_i}$ is over all neighboring cells $C_i$ of vertex $i$. Similarly, the tension of cell vertices to the cell center can be expressed as,

$$\Lambda^{(\mathrm{viscous})}_{i,J} = \eta^{(\mathrm{b})}_{J} \boldsymbol{t}_{i,J} \cdot (\boldsymbol{v}_J - \boldsymbol{v}_i)\,, \tag{S7}$$

where $\eta^{(\mathrm{b})}_{J}$ is the bulk viscous coefficient of the cell $J$; $\boldsymbol{t}_{i,J} = (\boldsymbol{r}_J - \boldsymbol{r}_i)/l_{i,J}$ with $l_{i,J} = |\boldsymbol{r}_i - \boldsymbol{r}_J|$ the distance of vertex $i$ to the geometric centre

$$\boldsymbol{r}_J = \frac{1}{n_J} \sum_{j \in cell} \boldsymbol{r}_j, \tag{S8}$$

of its neighbor cell $J$ with $n_J$ the number of vertices of cell $J$; $\boldsymbol{v}_J = \mathrm{d}\boldsymbol{r}_J/\mathrm{d}t$ is the velocity of the geometric centre of cell $J$. Since

$$\boldsymbol{v}_J = \frac{1}{n_J} \sum_{j \in cell\ J} \frac{\mathrm{d}\boldsymbol{r}_j}{\mathrm{d}t} = \frac{1}{n_J} \sum_{j \in cell\ J} \boldsymbol{v}_j, \tag{S9}$$

the viscous line tension $\Lambda^{(\mathrm{viscous})}_{i,J}$ between vertex $i$ and its neighboring cell $J$ can be further expressed as,

$$\Lambda^{(\mathrm{viscous})}_{i,J} = \eta^{(\mathrm{b})}_{J} \boldsymbol{t}_{i,J} \cdot (\boldsymbol{v}_J - \boldsymbol{v}_i) = \eta^{(\mathrm{b})}_{J} \boldsymbol{t}_{i,J} \cdot \left( \frac{1}{n_J} \sum_{j \in cell\ J} \boldsymbol{v}_j - \boldsymbol{v}_i \right) = \eta^{(\mathrm{b})}_{J} \boldsymbol{t}_{i,J} \cdot \left( -\frac{n_J - 1}{n_J} \boldsymbol{v}_i + \frac{1}{n_J} \sum_{\substack{j \in cell\ J \\ j \neq i}} \boldsymbol{v}_j \right). \tag{S10}$$

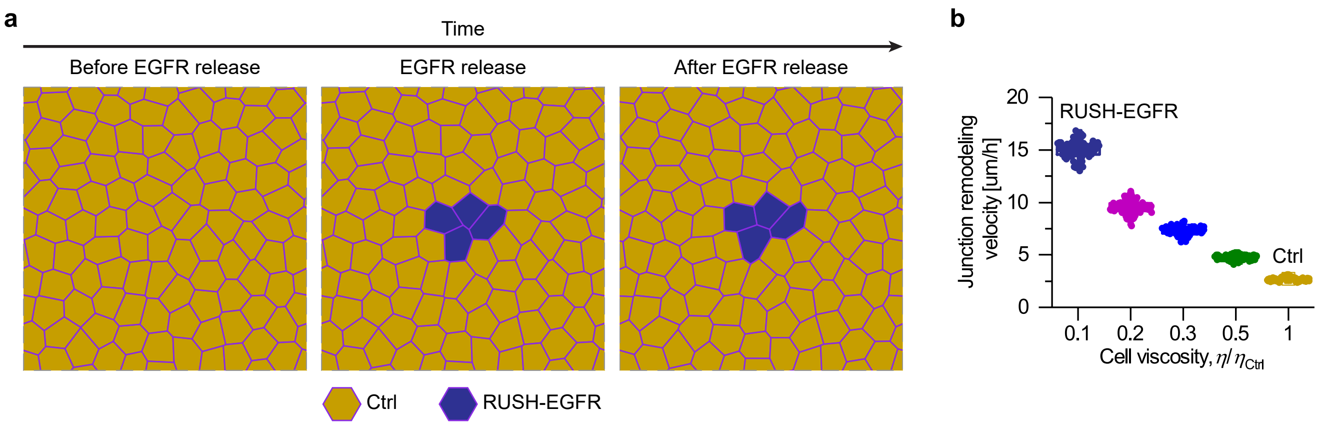


Supplementary Figure 2: Simulation of EGFR release experiments. (a) Model schematic. Before EGFR release, we simulate a cell sheet with active tension fluctuations. At steady state, we randomly select four contacting cells and decrease their cell viscosity from the Ctrl case, $\eta_{\rm Ctrl}$, to the RUSH-EGFR case, $\eta_{\rm RUSH-EGFR} = \eta_{\rm Ctrl}/10$. We then simulate the system and examine how the junction remodeling velocity changes. (b) The junction remodeling velocity as a function of the rescaled cell viscosity, $\eta/\eta_{\rm Ctrl}$ with $\eta_{\rm Ctrl}$ being the viscosity of control cells. One data point represents an average value of junction remodeling velocity of all junctions at a time frame ($n_{0.1} = 101$; $n_{0.2} = 101$; $n_{0.3} = 101$; $n_{0.5} = 101$; $n_1 = 101$). See Table I for default parameter values used in simulations.

Therefore, the viscous force Eq. (S4) can be expressed with respect to vertices' velocity $\{\boldsymbol{v}_i\}$ as,

$$\boldsymbol{F}_i^{\rm (viscous)} = \left[-\sum_{j\in V_i} \eta_{ij}^{\rm (s)} \boldsymbol{t}_{i,j} \otimes \boldsymbol{t}_{i,j} - \sum_{J\in C_i} \eta_J^{\rm (b)} \left(1 - \frac{1}{n_J}\right) \boldsymbol{t}_{i,J} \otimes \boldsymbol{t}_{i,J}\right] \cdot \boldsymbol{v}_i + \sum_{j\in V_i} \eta_{ij}^{\rm (s)} \left(\boldsymbol{t}_{i,j} \otimes \boldsymbol{t}_{i,j}\right) \cdot \boldsymbol{v}_j + \sum_{J\in C_i} \left(\eta_J^{\rm (b)} \frac{1}{n_J} \boldsymbol{t}_{i,J} \otimes \boldsymbol{t}_{i,J}\right) \cdot \sum_{\substack{j\,\in\, cell\\ j\neq i}} \boldsymbol{v}_j. \tag{S11}$$

##### C. Numerical scheme

**Initialization procedure** We initialize our simulations through a random Voronoi cellular pattern, using the procedure as described in our previous study [5].

**Motion procedure** Our procedure is the following: (i) Updating the active tension according to the Gillespie algorithm (see below); (ii) Solving the force balance equation to obtain the motion velocity of each vertex $\{\boldsymbol{v}_i\}$, using the method to be described in detail below; (iii) Move vertices to new positions using a forward Euler scheme,

$$\boldsymbol{r}_i(t + \Delta t) = \boldsymbol{r}_i + \boldsymbol{v}_i \Delta t; \tag{S12}$$

(iv) Perform T1 topological transitions for all short cell–cell junctions ($l_{ij} < \ell_{\rm T1} = 0.01\sqrt{A_0}$).

**Solving the force balance equation** To obtain the motion velocity $\boldsymbol{v}_i$ of each vertex $i$, we need to solve the force balance equation (S1). Substituting the expression of the friction force, Eq. (S2), and the viscous force, Eq. (S11), into the force balance equation (S1), we obtain

$$\left[\xi \boldsymbol{I} + \sum_{j\in V_i} \eta_{ij}^{\rm (s)} \left(\boldsymbol{t}_{i,j} \otimes \boldsymbol{t}_{i,j}\right) + \sum_{J\in C_i} \left(\eta_J^{\rm (b)} \frac{n_J - 1}{n_J} \boldsymbol{t}_{i,J} \otimes \boldsymbol{t}_{i,J}\right)\right] \cdot \boldsymbol{v}_i - \sum_{j\in V_i} \eta_{ij}^{\rm (s)} \left(\boldsymbol{t}_{i,j} \otimes \boldsymbol{t}_{i,j}\right) \cdot \boldsymbol{v}_j - \sum_{J\in C_i} \left(\eta_J^{\rm (b)} \frac{1}{n_J} \boldsymbol{t}_{i,J} \otimes \boldsymbol{t}_{i,J}\right) \cdot \sum_{\substack{j\in cell\ J\\ j\neq i}} \boldsymbol{v}_j = \boldsymbol{F}_i^{\rm (t)}, \tag{S13}$$

where

$$\boldsymbol{F}_i^{(\mathrm{t})} = \boldsymbol{F}_i^{(\mathrm{elastic})} + \boldsymbol{F}_i^{(\mathrm{active})}, \tag{S14}$$

is the total force that is independent of cell vertex velocity $\{\boldsymbol{v}_i\}$. The matrix form of Eq. (S13) reads,

$$\sum_{j=1}^{n} \boldsymbol{C}_{ij} \cdot \boldsymbol{v}_j = \boldsymbol{F}_i^{(\mathrm{t})}, \tag{S15}$$

where

$$\boldsymbol{C}_{ij} = \boldsymbol{C}_{ij}^{(\mathrm{f})} + \boldsymbol{C}_{ij}^{(\mathrm{s})} + \boldsymbol{C}_{ij}^{(\mathrm{b})}. \tag{S16}$$

Here $\boldsymbol{C}_{ij}^{(\mathrm{f})}$, $\boldsymbol{C}_{ij}^{(\mathrm{s})}$ and $\boldsymbol{C}_{ij}^{(\mathrm{b})}$ are the coefficient matrices corresponding to the cell–substrate friction, the cell-–cell interfacial viscosity and the cell bulk viscosity, with the detailed expression as below:

$$\boldsymbol{C}_{ij}^{(\mathrm{f})} = \xi \boldsymbol{I} \delta_{ij}, \tag{S17}$$

$$\boldsymbol{C}_{ij}^{(\mathrm{s})} = \begin{cases} \sum_{k \in V_i} \eta_{ik}^{(\mathrm{s})} \left(\boldsymbol{t}_{i,k} \otimes \boldsymbol{t}_{i,k}\right) & , \quad j = i \\ -\eta_{ij}^{(\mathrm{s})} \left(\boldsymbol{t}_{i,j} \otimes \boldsymbol{t}_{i,j}\right) & , \quad j \in V_i \\ \boldsymbol{0} & , \quad \text{otherwise} \end{cases} \tag{S18}$$

$$\boldsymbol{C}_{ij}^{(\mathrm{b})} = \begin{cases} \sum_{J \in C_i} \eta_J^{(\mathrm{b})} \frac{(n_J - 1)}{n_J} \left(\boldsymbol{t}_{i,J} \otimes \boldsymbol{t}_{i,J}\right) & , \quad j = i \\ -\sum_{i,j \in cell\ J} \eta_J^{(\mathrm{b})} \frac{1}{n_J} \boldsymbol{t}_{i,J} \otimes \boldsymbol{t}_{i,J} & , \quad j \neq i \\ \boldsymbol{0} & , \quad \text{otherwise} \end{cases} \tag{S19}$$

Equation (S15) gives the matrix form of the force balance at all vertices $i = 1, 2, \cdots, N$. We can further express all the force balance equations in a unified matrix form as below:

$$\boldsymbol{C} \cdot \boldsymbol{v} = \boldsymbol{F}^{(\mathrm{t})}, \tag{S20}$$

where $\boldsymbol{C}$ is referred to as the friction-viscosity coefficient matrix; $\boldsymbol{v}$ is the velocity vector of all vertices; $\boldsymbol{F}^{(\mathrm{t})}$ is the force vector of all vertices. $\boldsymbol{C}$, $\boldsymbol{v}$ and $\boldsymbol{F}^{(\mathrm{t})}$ are organized by $\boldsymbol{C}_{ij}$, $\boldsymbol{v}_i$ and $\boldsymbol{F}_i$ as below:

$$\boldsymbol{C} = \left(\boldsymbol{C}_{ij}\right)_{2N \times 2N} = \begin{pmatrix} \boldsymbol{C}_{11} & \ldots & \boldsymbol{C}_{1N} \\ \vdots & \ddots & \vdots \\ \boldsymbol{C}_{N1} & \cdots & \boldsymbol{C}_{NN} \end{pmatrix}_{2N \times 2N}, \tag{S21}$$

$$\boldsymbol{v} = \left(\boldsymbol{v}_i\right)_{2N \times 1} = \begin{pmatrix} \boldsymbol{v}_1 \\ \vdots \\ \boldsymbol{v}_N \end{pmatrix}_{2N \times 1}, \tag{S22}$$

$$\boldsymbol{F}^{(\mathrm{t})} = \left(\boldsymbol{F}_i^{(\mathrm{t})}\right) = \begin{pmatrix} \boldsymbol{F}_1^{(\mathrm{t})} \\ \vdots \\ \boldsymbol{F}_N^{(\mathrm{t})} \end{pmatrix}_{2N \times 1}, \tag{S23}$$

where $N$ is the total number of cell vertices. Solving the system of linear algebraic equations (S20) together with the boundary conditions gives the motion velocity of each vertex $\{\boldsymbol{v}_i\}$.

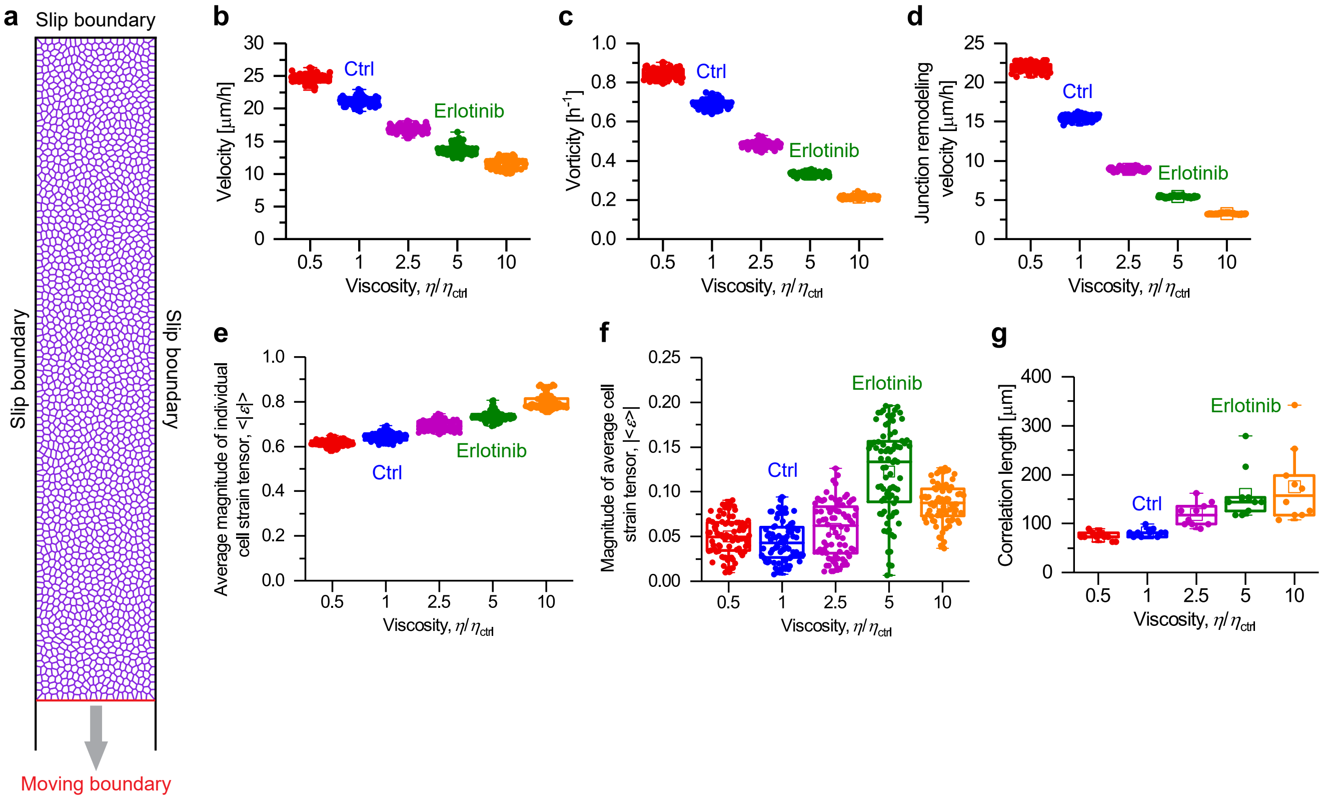


Supplementary Figure 3: Simulation of collective cell migration experiments. (a) Model schematic: tissue geometry and boundary conditions. (b-e) Cell velocity (b), cell vorticity (c), junction remodeling velocity (d), average magnitude of individual cell strain tensor (e), magnitude of average cell strain tensor (f), and correlation length (g) as a function of the viscosity. One data point represents an average value of junction remodeling velocity of all junctions at a time frame: in (b-f), $n_{0.5} = 80$, $n_1 = 80$, $n_{2.5} = 80$, $n_5 = 80$, $n_{10} = 80$; in (g), $n_{0.5} = 10$, $n_1 = 10$, $n_{2.5} = 10$, $n_5 = 10$, $n_{10} = 10$. See Table II for default parameter values used in simulations.

**Topological transitions** We implement the T1 topological transition to mimic cell intercalation as observed in experiments. A T1 topological transition is performed once a cell–cell interface becomes shorter than a threshold (i.e., $l_{i,j} < \ell_{\mathrm{T1}}$) and accepted only when such a topological transition lowers the mechanical energy of the cell monolayer system (Eq. (S3)). In detail, a T1 topological transition is accomplished by rotating the corresponding cell–cell interface by 90 degrees and updating the vertices lists of the four cells involved.

**Updating active tensions (RUSH-EGFR model)** Active tension fluctuations contribute to the following force at each vertex,

$$\boldsymbol{F}_i^{(\mathrm{active})} = \sum_{j \in \mathrm{neighbor}} \Lambda_{ij}^{(\mathrm{act})} \boldsymbol{t}_{i,j}, \tag{S24}$$

where the summation is over all neighbor vertices that connect to the vertex $i$. Numerically, we update the active tension at all cell-cell junctions using the Gillespie algorithm [7, 8], that is,

$$\Lambda_{ij}^{(\mathrm{act})}(t+\Delta t) = \Lambda_{ij}^{(\mathrm{act})}(t)\exp\left(-\frac{\Delta t}{\tau_\sigma}\right) + \Delta_\sigma \sqrt{\frac{\tau_\sigma}{2}\left[1-\exp\left(-\frac{2\Delta t}{\tau_\sigma}\right)\right]}\,\upsilon_{ij}(t), \tag{S25}$$

where $\upsilon_{ij}(t)$ is a random number that satisfies the normal distribution, $\upsilon_{ij}(t) \sim \mathcal{N}(0,1)$.

**Cell migration experiments** The cell migration experiment method is detailed in the main text, Method section. We estimated several key quantities for a set of viscosity ratios, see Supplementary Figs. 3 and 4.

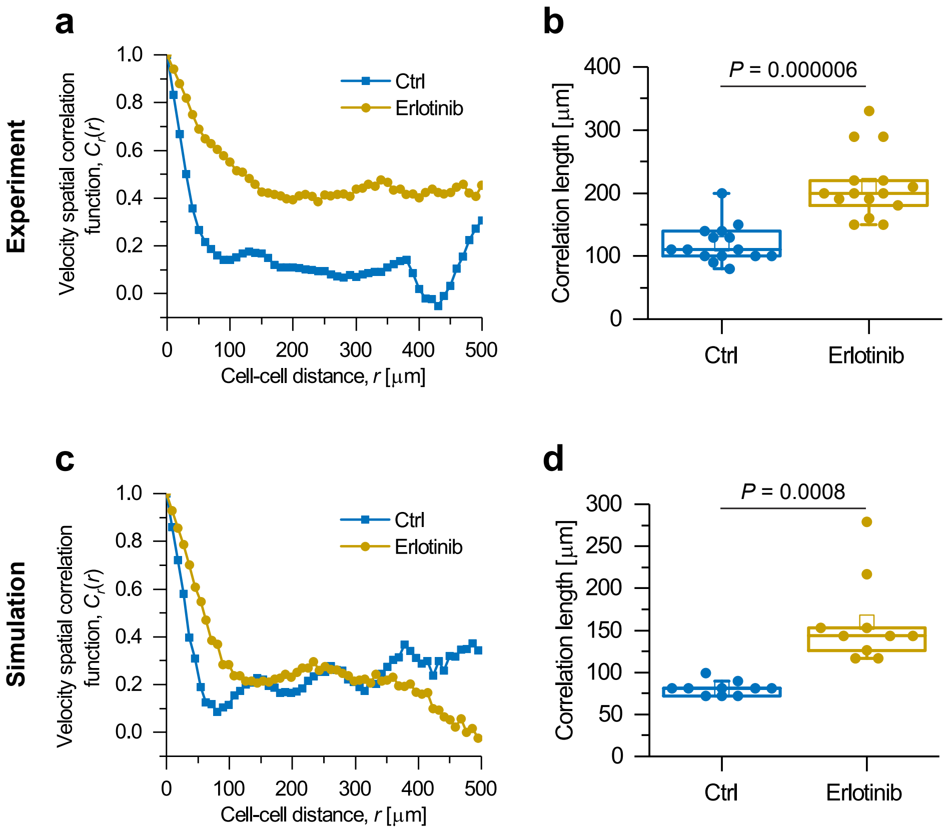


Supplementary Figure 4: Simulation of collective cell migration experiments. Comparison of (a, c) the velocity spatial correlation function $C_r(r)$ and (b, d) the correlation length between (a, b) experiments and (c, d) simulations. (a, c) Shown here are the typical velocity spatial correlation function curves for (a) experiments and (c) simulations. (b, d) Comparison of the correlation length extracted from the velocity spatial correlation function (see main text, Method section). Each data point represents a correlation length value evaluated at a time frame: for experiments, $n_{\mathrm{Ctrl}} = n_{\mathrm{Erlotinib}} = 15$; for simulations, $n_{\mathrm{Ctrl}} = n_{\mathrm{Erlotinib}} = 10$. $\boldsymbol{P}$ values were calculated using Student's two-tailed unpaired $t$-tests. See Table II for default parameter values used in simulations.

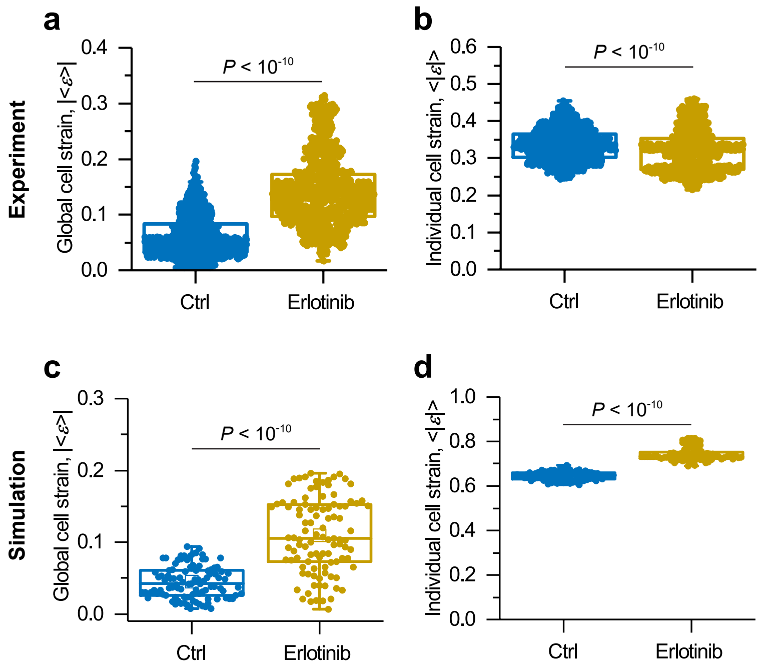


Supplementary Figure 5: Simulation of collective cell migration experiments. Comparison of (a, c) the global cell strain $|\langle \varepsilon \rangle|$ and (b, d) the individual cell strain $\langle |\varepsilon| \rangle$ between (a, b) experiments and (c, d) simulations. Here, $\langle \cdot \rangle$ means an average over space. Each data point represents a value evaluated at a time frame: for experiments, $n_{\mathrm{Ctrl}} = n_{\mathrm{Erlotinib}} = 1060$; for simulations, $n_{\mathrm{Ctrl}} = n_{\mathrm{Erlotinib}} = 100$. $\boldsymbol{P}$ values were calculated using Student's two-tailed unpaired $t$-tests. See Table II for default parameter values used in simulations.

##### D. Evaluation of tissue viscoelastic relaxation time

###### 1. Tensor maps

**Individual cell strain tensor** We quantify the cell strain based on the texture tensor measurement as [9, 10],

$$\boldsymbol{\varepsilon}_J = \frac{1}{2}\ln\left(\frac{\boldsymbol{M}_J}{\boldsymbol{M}_0}\right), \tag{S26}$$

where $\boldsymbol{M}_J$ is the texture tensor of the $J$-th cell, defined as

$$\boldsymbol{M}_J = \frac{1}{N_J}\sum_{K\in neighbor}\left(\boldsymbol{r}_K - \boldsymbol{r}_J\right)\otimes\left(\boldsymbol{r}_K - \boldsymbol{r}_J\right), \tag{S27}$$

with $N_J$ the number of neighbor cells to the cell $J$; $\boldsymbol{M}_0$ represents a reference texture tensor [9]. We define $\boldsymbol{M}_0$ as the texture tensor of a regular hexagonal cell pattern in a stress-free state [5].

**Texture field $\boldsymbol{M}$** In all that follows, we will define coarse-grained tensors ($\boldsymbol{M}$, $\boldsymbol{\varepsilon}_{\text{cell}}$, $\boldsymbol{v}$), each of them being evaluated on a set of points of a regular grid. Following [9], we define the texture tensor $\boldsymbol{M}$ through the averaging procedure

$$\boldsymbol{M}\left(\boldsymbol{r}\right) = \langle \boldsymbol{M}_J\rangle = \frac{\sum\limits_{|\boldsymbol{r}-\boldsymbol{r}_J|<r_{\text{cut-off}}} w\left(\boldsymbol{r}-\boldsymbol{r}_J\right)\boldsymbol{M}_J}{\sum\limits_{|\boldsymbol{r}-\boldsymbol{r}_J|<r_{\text{cut-off}}} w\left(\boldsymbol{r}-\boldsymbol{r}_J\right)}, \tag{S28}$$

where $\boldsymbol{r}_J = \sum_{j\in cell}\boldsymbol{r}_j/n_J$ is the cell geometric centre; where $w(\boldsymbol{r}-\boldsymbol{r}_J)$ is the weight function and $r_{\text{cut}-\text{off}} = 3\sigma$ is a cut-off length. Here we set the weight function as a Gaussian function,

$$w\left(\boldsymbol{r}-\boldsymbol{r}_J\right) = \frac{1}{\sqrt{2\pi}\sigma}\exp\left(-\frac{1}{2}\frac{\left|\boldsymbol{r}-\boldsymbol{r}_J\right|^2}{\sigma^2}\right), \tag{S29}$$

where we set the kernel size $\sigma$ at $\sigma = 0.75$ cell length. We set the cut-off length at $r_{\text{cut}-\text{off}} = 3\sigma = 2.25$ (in the cell length unit).

**Strain tensor and velocity vector fields** Based on the strain tensor ($\boldsymbol{\varepsilon}_J$),following the same method as $\boldsymbol{M}$, we define a coarse-grained cell strain field

$$\boldsymbol{\varepsilon}_{\text{cell}} = \langle\boldsymbol{\varepsilon}_J\rangle. \tag{S30}$$

Similarly, we construct the velocity field $\boldsymbol{v}$ based on the individual cell velocity vector $\boldsymbol{v}_J$. Further, based on the coarse-grained velocity field, we calculate the strain rate tensor $\nabla\boldsymbol{v}$ via the finite difference method.

**Topological texture change tensor $\boldsymbol{T}$** Following [9], we define the topological texture change tensor due to cell rearrangements defined as [9],

$$\boldsymbol{T} = \dot{n}_a\langle\boldsymbol{\ell}_a\otimes\boldsymbol{\ell}_a\rangle - \dot{n}_d\langle\boldsymbol{\ell}_d\otimes\boldsymbol{\ell}_d\rangle, \tag{S31}$$

where $\langle\rangle$ is the averaging procedure defined in Eq. (S28); $\boldsymbol{\ell}_a$ and $\boldsymbol{\ell}_d$ are appearing cell–cell links and disappearing cell–cell links, respectively; the quantity $\dot{n}_a$ (respectively, $\dot{n}_d$) refers to the rate of cell–cell link appearance (respectively, disappearance), per unit time and per existing cell–cell link. Computationally, we evaluate the quantity:

$$\boldsymbol{T}(\boldsymbol{r}) = \frac{N_{\text{T1}}}{\Delta T N_{\text{link}}}\frac{1}{\sum\limits_{j=1,\ldots,N_{\text{T1}}} w\left(\boldsymbol{r}-\boldsymbol{r}_j\right)}\sum_{j=1,\ldots,N_{\text{T1}}} w\left(\boldsymbol{r}-\boldsymbol{r}_j\right)\left(\boldsymbol{\ell}_a^j\otimes\boldsymbol{\ell}_a^j - \boldsymbol{\ell}_d^j\otimes\boldsymbol{\ell}_d^j\right), \tag{S32}$$

where $N_{\text{T1}}$ is the number of rearrangements that occurred during the time duration $\Delta T$ within the box centered on $\boldsymbol{r}$ and of size $r_{\text{cut-off}}$; $N_{\text{link}}$ is the average number of cell-cell links within the box centered on $\boldsymbol{r}$ and of size $r_{\text{cut-off}}$.

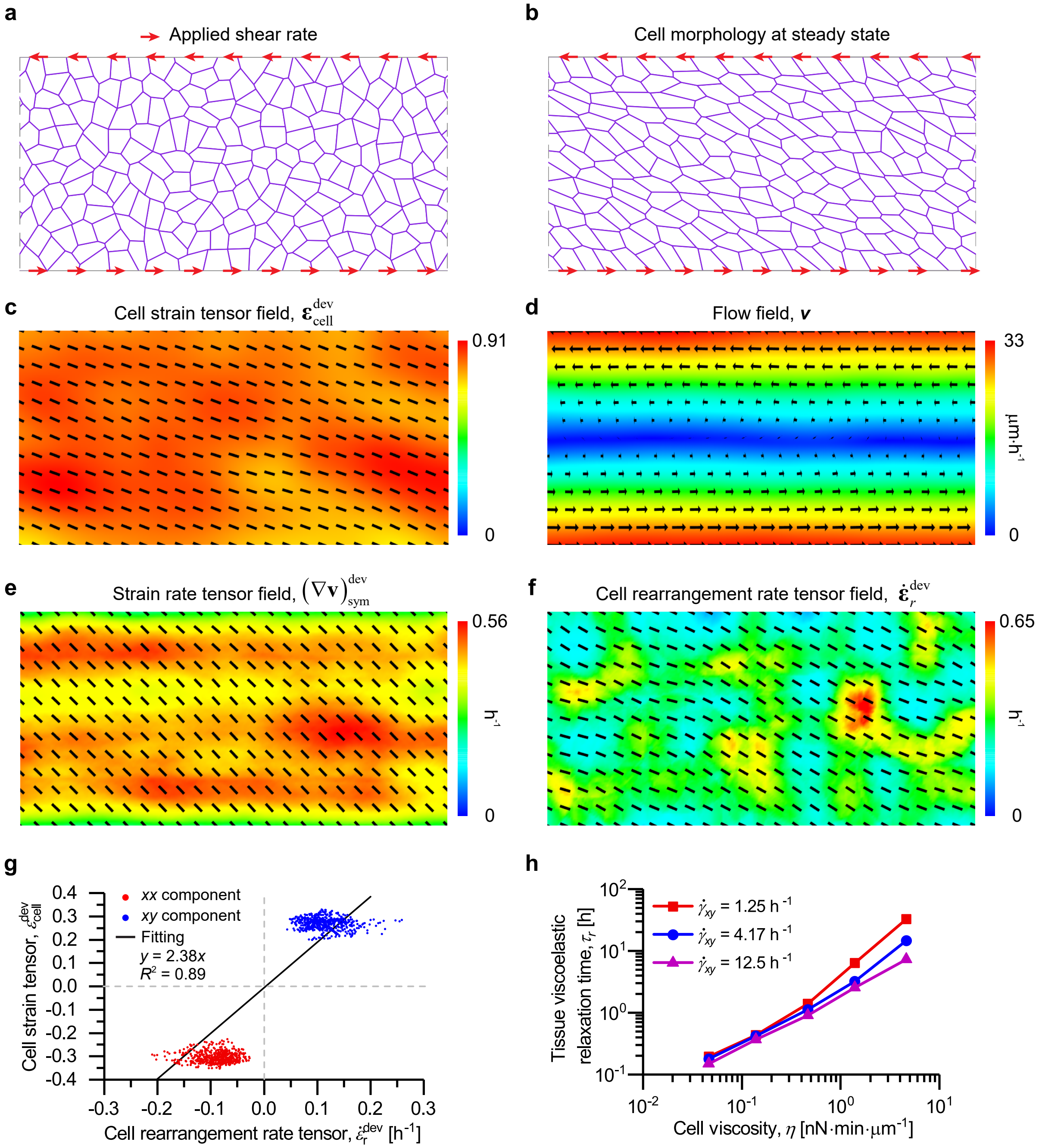


Supplementary Figure 6: Evaluation of tissue viscoelastic relaxation time. (a) Model schematic. We apply a constant shear rate at the top and bottom boundary vertices. (b) Cell morphology at steady state. (c) The cell deformation strain field, $\boldsymbol{\varepsilon}_{\mathrm{cell}}^{\mathrm{dev}}$ (the deviatoric part), where the color code refers to the magnitude and arrows represent orientations. Averaged over $N_{\mathrm{frame}} = 500$ frames at steady state. (d) The flow field, where the color code refers to the velocity magnitude and arrows represent velocity vectors. Averaged over $N_{\mathrm{frame}} = 500$ frames at steady state. (e) The strain rate tensor field, $(\nabla \boldsymbol{v})_{\mathrm{sym}}^{\mathrm{dev}}$ (the deviatoric part), where the color code refers to the magnitude and arrows represent orientations. Averaged over $N_{\mathrm{frame}} = 500$ frames at steady state. (f) The cell rearrangement rate tensor field, $\dot{\boldsymbol{\varepsilon}}_r^{\mathrm{dev}}$ (the deviatoric part), where the color code refers to the magnitude and arrows represent orientations. Averaged over $N_{\mathrm{frame}} = 500$ frames at steady state. (g) Fitting the relaxation time via equation (S35), based on the data of cell rearrangement rate tensor, $\dot{\boldsymbol{\varepsilon}}_{\mathrm{r}}^{\mathrm{dev}}$, and cell strain tensor, $\boldsymbol{\varepsilon}_{\mathrm{cell}}^{\mathrm{dev}}$. (h) Tissue viscoelastic relaxation time $\tau_r$ as a function of cell viscosity $\eta$, at different shear rates $\dot{\gamma}_{xy}$. In (b-g), $\dot{\gamma}_{xy} = 0.42\ \mathrm{h}^{-1}$, $\eta = 1.4\ \mathrm{nN} \cdot \mathrm{min} \cdot \mu\mathrm{m}^{-1}$. Parameters: $\xi = 0$; see Table I for other default parameter values used in simulations.

**Cell rearrangement rate tensor** From $\boldsymbol{M}$ and $\boldsymbol{T}$, defined in Eqs. (S28) and (S31), we quantity the cell rearrangement rate tensor field based on T1 topological transition events as [9],

$$\dot{\boldsymbol{\varepsilon}}_r = -\frac{1}{2}\big(\boldsymbol{M}^{-1} \cdot \boldsymbol{T}\big)_{\rm sym}, \tag{S33}$$

where $()_{\rm sym}$ is a tensor symmetrization operator.

##### 2. *Extraction of a viscoelastic relaxation time*

Here we detail how we perform shear simulations to evaluate the tissue viscoelastic relaxation time.

We simulate a cell sheet consisting of $N = 200$ cells in a slab geometry of size $L_x = 20\sqrt{A_0}$ and $L_y = 10\sqrt{A_0}$, see Supplementary Figure 6a. To mimic a constant applied shear rate, we apply a constant velocity at the top ($y = +L_y/2$) and bottom ($y = -L_y/2$) vertices,

$$\boldsymbol{v}_{\rm boundary} = y\dot{\gamma}_{xy}\hat{\boldsymbol{x}}, \quad \text{for } y = \pm\frac{L_y}{2} \tag{S34}$$

Here, $\dot{\gamma}_{xy}$ is the applied shear rate parameter. Such boundary conditions are implemented into the force balance equation (S20) using a Lagrange multiplier method. In addition, we assume periodic boundary conditions along the $x$ axis.

Due to the applied shear rate, the simulated tissue exhibits a well-oriented cellular pattern with large cell deformation (Supplementary Figure 6b,c), a well-defined shear flow pattern (Supplementary Figure 6d), a well-defined strain rate pattern $\boldsymbol{\varepsilon}_{\rm cell}$ as defined per Eq. (S30) (Supplementary Figure 6e), and cell rearrangement rate tensor field, $\dot{\boldsymbol{\varepsilon}}_r$ as defined per Eq. (S33) (Supplementary Figure 6f).

We then verified the following linear relation between the strain and cell rearrangement tensors

$$\dot{\boldsymbol{\varepsilon}}_r = \frac{\boldsymbol{\varepsilon}_{\rm cell}}{\tau_r}, \tag{S35}$$

which defines a characteristic time constant $\tau_r$. Such linear relation is consistent with a Maxwell viscoelastic model, hence $\tau_r$ is referred to the a viscoelastic constant [11]. We provide an example of simulation data fit in Supplementary Figure 6g.

TABLE I: List of default parameter values used in simulations of EGFR release experiments.

| Parameter | Description | Value | Note |
|---|---|---|---|
| $\ell = \sqrt{A_0}$ | Length scale | 18 $\mu$m | Estimated |
| $\tau$ | Time scale | 0.15 min | Estimated |
| $f = K_A A_0^{3/2}$ | Force scale | 5.8 nN | Estimated |
| $K_A$ | Cell area stiffness | $10^6$ N $\cdot$ m$^{-3}$ | ref. [12] |
| $A_0$ | Preferred cell area | 324 $\mu$m$^2$ | Estimated |
| $K_P$ | Cell perimeter stiffness | $6.5 \times 10^{-3}$ nN $\cdot$ $\mu$m$^{-1}$ | ref. [13] |
| $P_0$ | Preferred cell perimeter | 18 $\mu$m | Assumed |
| $\xi$ | Cell-substrate friction | 0.28 nN $\cdot$ min $\cdot$ $\mu$m$^{-1}$ | Assumed |
| $\Delta_\sigma$ | Magnitude of active tension fluctuations | 0.15 nN $\cdot$ min$^{-1/2}$ | Assumed |
| $\tau_\sigma$ | Correlation time of active tension fluctuations | 1.5 min | Assumed |
| $\eta_{\mathrm{Ctrl}}$ | Viscosity of control cells | 1.4 nN $\cdot$ min $\cdot$ $\mu$m$^{-1}$ | Estimated |
| $\eta_{\mathrm{RUSH-EGFR}}$ | Viscosity of RUSH-EGFR cells | 0.14 nN $\cdot$ min $\cdot$ $\mu$m$^{-1}$ | Estimated |
| $\Delta t$ | Simulation time step | 0.015 min | Assumed |

TABLE II: List of default parameter values used in simulations of cell migration experiments.

| Parameter | Description | Value | Note |
|---|---|---|---|
| $\ell = \sqrt{A_0}$ | Length scale | 18 $\mu$m | Estimated |
| $\tau$ | Time scale | 0.15 min | Estimated |
| $f = K_A A_0^{3/2}$ | Force scale | 5.8 nN | Estimated |
| $K_A$ | Cell area stiffness | $10^6$ N $\cdot$ m$^{-3}$ | ref. [12] |
| $K_P$ | Cell perimeter stiffness | $6.5 \times 10^{-3}$ nN $\cdot$ $\mu$m$^{-1}$ | ref. [13] |
| $P_0$ | Preferred cell perimeter | 18 $\mu$m | Assumed |
| $\xi$ | Cell-substrate friction | 0.28 nN $\cdot$ min $\cdot$ $\mu$m$^{-1}$ | Assumed |
| $T_0$ | Traction force magnitude | 0.58 nN | ref. [14] |
| $V_{\mathrm{front}}$ | Moving front velocity | 15 $\mu$m $\cdot$ h$^{-1}$ | Estimated |
| $\eta_{\mathrm{Erlotinib}}$ | Viscosity of EGFR-inhibited cells | 1.4 nN $\cdot$ min $\cdot$ $\mu$m$^{-1}$ | Estimated |
| $\eta_{\mathrm{Ctrl}}$ | Viscosity of control cells | 0.28 nN $\cdot$ min $\cdot$ $\mu$m$^{-1}$ | Estimated |
| $\Delta t$ | Simulation time step | 0.015 min | Assumed |

[1] T. Nagai and H. Honda, Philosophical Magazine B **81**, 699 (2001).
[2] R. Farhadifar, J. C. Röper, B. Algouy, S. Eaton, and F. Jülicher, Current Biology **17**, 2095–2104 (2007).
[3] S. Alt, P. Ganguly, and G. Salbreux, Philosophical Transactions of the Royal Society B: Biological Sciences **372**, 20150520 (2017).
[4] A. G. Fletcher, M. Osterfield, R. E. Baker, and S. Y. Shvartsman, Biophysical Journal **106**, 2291–2304 (2014).
[5] S.-Z. Lin, M. Merkel, and J.-F. Rupprecht, Physical Review Letters **130**, 058202 (2023).
[6] S.-Z. Lin, S. Ye, G.-K. Xu, B. Li, and X.-Q. Feng, Biophysical Journal **115**, 1826 (2018).
[7] D. T. Gillespie, Physical Review E **54**, 2084 (1996).
[8] S. Tlili, J. Yin, J.-F. Rupprecht, M. A. Mendieta-Serrano, G. Weissbart, N. Verma, X. Teng, Y. Toyama, J. Prost, and T. E. Saunders, Proceedings of the National Academy of Sciences of the United States of America **116**, 25430–25439 (2019).
[9] F. Graner, B. Dollet, C. Raufaste, and P. Marmottant, The European Physical Journal E **25**, 349 (2008).
[10] S. Tlili, C. Gay, F. Graner, P. Marcq, F. Molino, and P. Saramito, The European Physical Journal E **38**, 1 (2015).
[11] S. Tlili, M. Durande, C. Gay, B. Ladoux, F. Graner, and H. Delanoë-Ayari, Physical Review Letters **125**, 088102 (2020).
[12] P. P. Girard, E. A. Cavalcanti-Adam, R. Kemkemer, and J. P. Spatz, Soft Matter **3**, 307 (2007).
[13] E. Hannezo, J. Prost, and J.-F. Joanny, Proceedings of the National Academy of Sciences of the United States of America **111**, 27 (2014).
[14] O. Cochet-Escartin, J. Ranft, P. Silberzan, and P. Marcq, Biophysical Journal **106**, 65 (2014).
